# Sleep renormalizes learning-perturbed cortical dynamics to stabilize memory

**DOI:** 10.64898/2026.03.16.711029

**Authors:** Thea Ng, Morgan Barnes, Atif Abedeen, Lucas Collignon, Hirni Patel, Nicholas Vovcsko, Rebecca M. C. Spencer

## Abstract

The brain must preserve learning-induced circuit changes without destabilizing the network dynamics required for future learning. We show that sleep addresses this tension through perturbation-dependent renormalization of cortical dynamics. Using high-density electroencephalography in humans following declarative learning and a matched non-learning control, we found that learning displaced cortical population dynamics during wakefulness, followed by an opposing reorganization during non-rapid eye movement (NREM) sleep. The preceding waking perturbation constrained this sleep response across time, cortical space, and individuals: NREM activity evolved along the wake-defined state-space direction, regions with larger waking perturbations showed stronger opposing responses, and larger perturbations predicted stronger restorative trajectories. This cross-state geometry generalized to held-out participants and predicted sleep-specific memory benefit beyond either state alone or their simple difference. Neural stability is therefore achieved by regulating trajectories rather than returning to a fixed baseline, with plasticity-induced perturbations specifying how cortical state space is reorganized to preserve learned structure while enabling further change.

## Introduction

Adaptive neural systems face a fundamental problem: experience must change the brain without destabilizing the dynamics that make further learning possible^1,2^. Memory compounds this problem because transforming initially labile representations into durable memories requires learning-induced circuit modifications, yet their persistence must remain compatible with on-going computation and future plasticity^3–5^. The challenge is therefore not simply to reverse the effects of plasticity, but to preserve what plasticity encodes without allowing the accompanying dynamical perturbation to persist. Sleep has long been proposed to reconcile these competing demands, stabilizing newly acquired information while restoring the capacity for subsequent learning^6–8^. At the synaptic level, the synaptic homeostasis hypothesis proposes that wake-associated increases in synaptic strength are renormalized during sleep to counteract saturation^6,9–12^. Whether sleep similarly counteracts learning-induced perturbations at the level of cortical population dynamics remains unclear. Current models of sleep-dependent consolidation have largely focused on how newly encoded memories are reactivated during non-rapid eye movement (NREM) sleep, particularly through coordinated interactions among slow oscillations, spindles and hippocampal ripples that facilitate communication across brain regions^13–15^. This work has revealed how memories are processed during sleep, but it leaves open how prior learning reshapes the broader population dynamics expressed across wake and sleep. More generally, a fundamental function of sleep may be to renormalize the cortical operating regime perturbed by learning, preserving learning-related structure while restoring the dynamic range required for subsequent adaptation.

If such restoration contributes to memory stabilization, a simple difference between wakefulness and sleep is not enough: neural activity changes substantially with vigilance state and elapsed time even in the absence of learning^16,17^. Homeostatic regulation across neural systems provides a powerful precedent: deviations from regulated neural activity recruit compensatory dynamics that oppose the preceding disturbance and preserve stable circuit function over time^2,18,19^. For example, chronically elevating activity in cortical networks initially increases firing, but compensatory weakening of excitatory synapses subsequently returns firing toward control levels^20^. Therefore, perturbation dependence should be a defining feature of restoration: the response should be organized by the disturbance it acts to correct, not by a single scalar endpoint. In other words, the response should retain information about what was perturbed. Applied to learning, this predicts that the magnitude and pattern of the waking perturbation should constrain subsequent sleep dynamics, defining the population dimension along which the system reorganizes. Averaged across the population, sleep should oppose the learning-induced shift established during wakefulness. Across states and over time within each state, population activity should evolve continuously, and the direction of this evolution should depend on the preceding perturbation. Thus, the relevant signature is the relation between successive operating regimes, rather than the level reached in either state alone. Finally, if this cross-state reorganization contributes to consolidation, the relationship between the learning-related waking perturbation and its subsequent transformation across sleep should predict the memory benefit of sleep.

The same principle should apply spatially. Learning-induced perturbations are unlikely to be distributed uniformly across cortex. Evidence from local sleep supports this possibility. Learning or experimentally induced cortical potentiation can increase subsequent slow-wave activity (SWA) over the engaged region, whereas immobilizing an arm produces a local decrease over the corresponding sensorimotor cortex^21–23^. But localization is not the same as spatial correspondence. Perturbation-dependent renormalization makes the stronger prediction that regional variation in the waking disturbance should predict regional variation in the subsequent compensatory response. For declarative learning, we expected this opposing wake–sleep organization to be most prominent over frontocentral cortex^24–26^. In addition, spatial correspondence captures only one part of this reorganization. Neural computations are increasingly understood in terms of the geometry and dimensionality of population activity, including the states populations occupy and the trajectories they form through state space^27–30^. This matters because the same average change can mask very different population dynamics. If restoration preserves the structure of the preceding perturbation, learning should shape not only where population activity lies in state space but the trajectory by which it subsequently reorganizes during sleep.

Testing this across wakefulness and sleep requires a signal that can be tracked across both states despite substantial differences in their population synchrony and oscillatory structure^16,31,32^. The aperiodic, 1/f-like component of neural power spectra provides such a coordinate. Its spectral slope varies with arousal and shows modulation with attention, perception and working-memory demands, with individual differences also related to cognitive performance^16,33–38^. In addition, REM-related aperiodic recalibration has previously been associated with overnight memory retention^39^. Although not a direct measure of any one cellular mechanism^40,41^, spectral slope is sensitive to properties that shape population activity over time, including excitation-inhibition balance and temporal correlation structure, and can provide a population-level readout of the operating regime in which neural activity is expressed^42–45^. A steeper spectrum places relatively more power at lower than higher frequencies, whereas a flatter spectrum does the reverse. This matters for sleep because band-limited power inherits the broadband structure of the spectrum. An increase in SWA, for example, can reflect a steeper slope, additional low-frequency activity, or both. Spectral parameterization distinguishes these contributions analytically without assuming that they arise from separate biological processes^46,47^. It also allows band-limited correlates of sleep homeostasis, including waking theta, NREM SWA and spindle-range sigma activity^25,26^, to be examined independently of aperiodic spectral changes. Broadband spectral organization may therefore reflect the operating regime within which frequency-specific sleep activity occurs^31,46,48^.

To test whether sleep renormalizes a cortical operating regime perturbed by learning, we recorded high-density EEG across a continuous wake–sleep cycle in humans following declarative word-pair learning and a matched non-learning control. We used aperiodic slope as a common cross-state coordinate to ask whether changes across the wake–sleep transition reflected brain state alone or were organized by the learning perturbation that preceded sleep. We show that learning displaces the cortical operating regime during wakefulness and that subsequent NREM activity follows a perturbation-dependent population trajectory opposing this shift. This cross-state population pattern generalizes to held-out participants, and its strength predicts the sleep-specific benefit to memory. These findings identify perturbation-dependent renormalization as a systems-level principle that may bring homeostatic regulation and consolidation into a common framework, suggesting that neural adaptability depends on coordinating learning-induced change with the subsequent reorganization of population dynamics.

## Results

To track how learning-related cortical population dynamics evolve from wakefulness into sleep, 36 healthy young adults completed a declarative word-pair paradigm together with high-density EEG recordings across Learning and Sham nights (Fig. 1a–c). Memory was tested immediately after encoding and again after either a wake or overnight-sleep interval to quantify retention. EEG was recorded during post-task wakeful rest and subsequent overnight sleep, and spectral parameterization separated aperiodic activity from oscillatory components, providing a common measure of cortical dynamics across brain states (Fig. 1d,e). Participant demographics and sleep characteristics are reported in Table S1.1.

**Figure 1:**
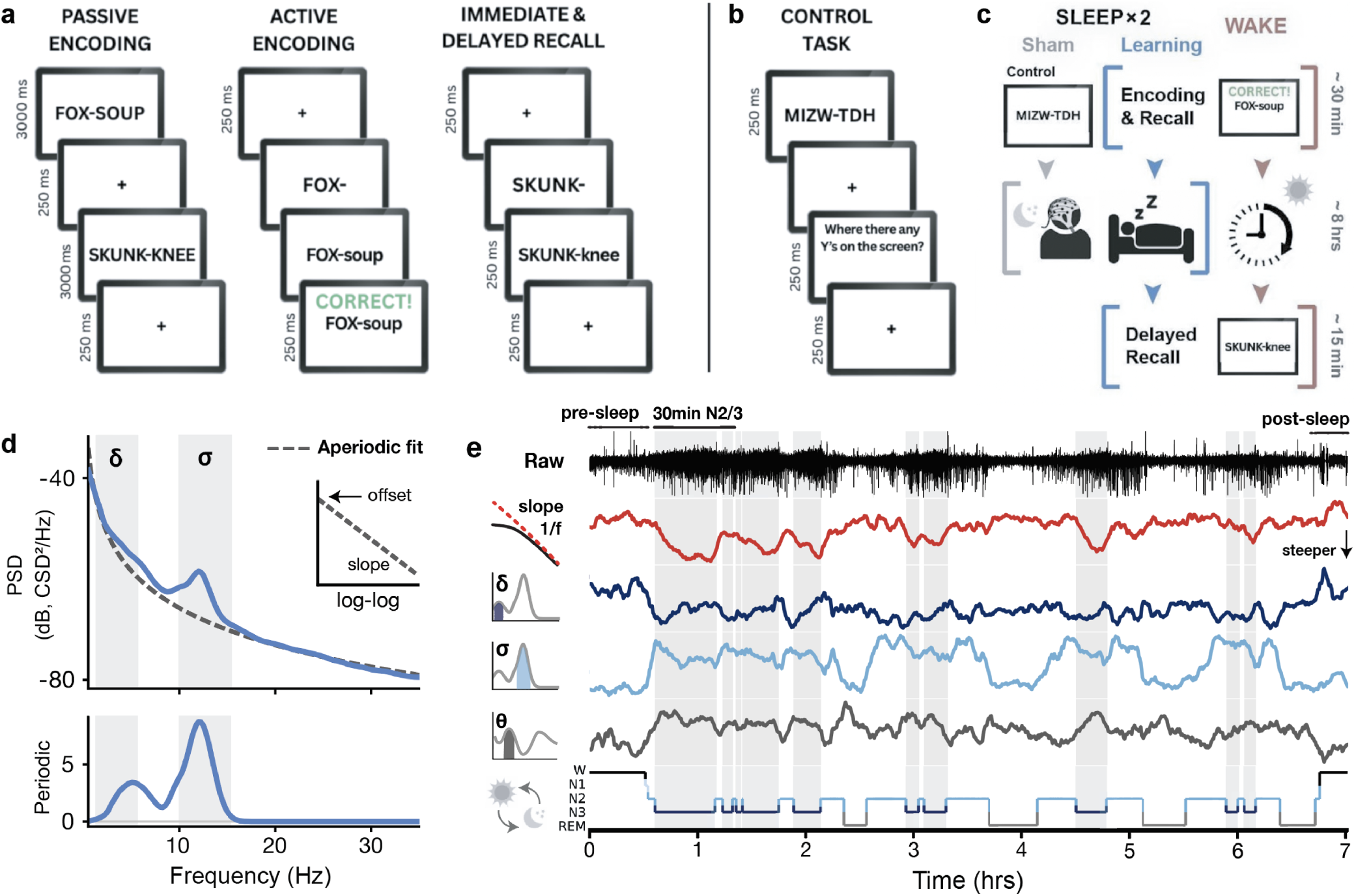
Experimental paradigm and spectral parameterization of cortical dynamics across wake and sleep. **a**, Declarative word-pair learning task. Thirty-six participants passively encoded 32 unrelated word pairs and completed repeated active cued-recall blocks with feedback until reaching criterion (≥20 correct responses within a block) or completing five blocks. Immediate and delayed recall tested the same word pairs without feedback. **b**, Sham task. Participants monitored pseudoword pairs for specified letters to match attentional demands without declarative learning or subsequent recall. **c**, Within-participant study design. Participants completed Wake, Sleep and Sham sessions on Days 7, 14 and 21. Sleep and Sham sessions were performed before bedtime and followed by overnight high-density polysomnography (PSG); delayed recall was completed the following morning after Sleep, whereas participants departed after awakening in the Sham session. The Wake session began in the morning and was followed by an 8-h waking retention interval without napping, before delayed recall in the evening. **d**, The observed power spectral density (PSD) is decomposed into an aperiodic 1/f component and superimposed oscillatory activity; the inset shows the aperiodic offset and slope in log–log coordinates, and the lower panel shows the resulting aperiodic-adjusted periodic spectrum. **e**, Representative overnight continuous trajectory at Fz after current source density (CSD) transformation. From top to bottom, traces show the CSD-transformed EEG signal, aperiodic 1/f slope, aperiodic-adjusted SWA (0.5–4 Hz), aperiodic-adjusted sigma power (10–16 Hz), aperiodic-adjusted pre-sleep theta power (4–8 Hz) and the hypnogram (Wake, N1, N2, N3 and REM). Aperiodic slope was estimated from 1–35-Hz *specparam* fits, and oscillatory measures from the aperiodic-adjusted residual spectrum. Light-gray shading indicates N3 sleep.

### Sleep preserves newly learned declarative memory

We first established the behavioral benefit of sleep relative to wakefulness for newly learned declarative memory. Participants varied substantially in the number of encoding blocks completed, and those requiring more blocks showed poorer accuracy on their final encoding block (Fig. 2a,b). Despite this variability, recall diverged across the subsequent retention interval. A binomial generalized linear mixed-effects model (GLMM) showed that accuracy declined from immediate to delayed testing across wakefulness but remained stable across overnight sleep. The decline was significantly greater across wakefulness than across sleep (Fig. 2c,d). This sleep-dependent preservation remained robust after accounting for encoding ability, sleepiness and word set.

**Figure 2:**
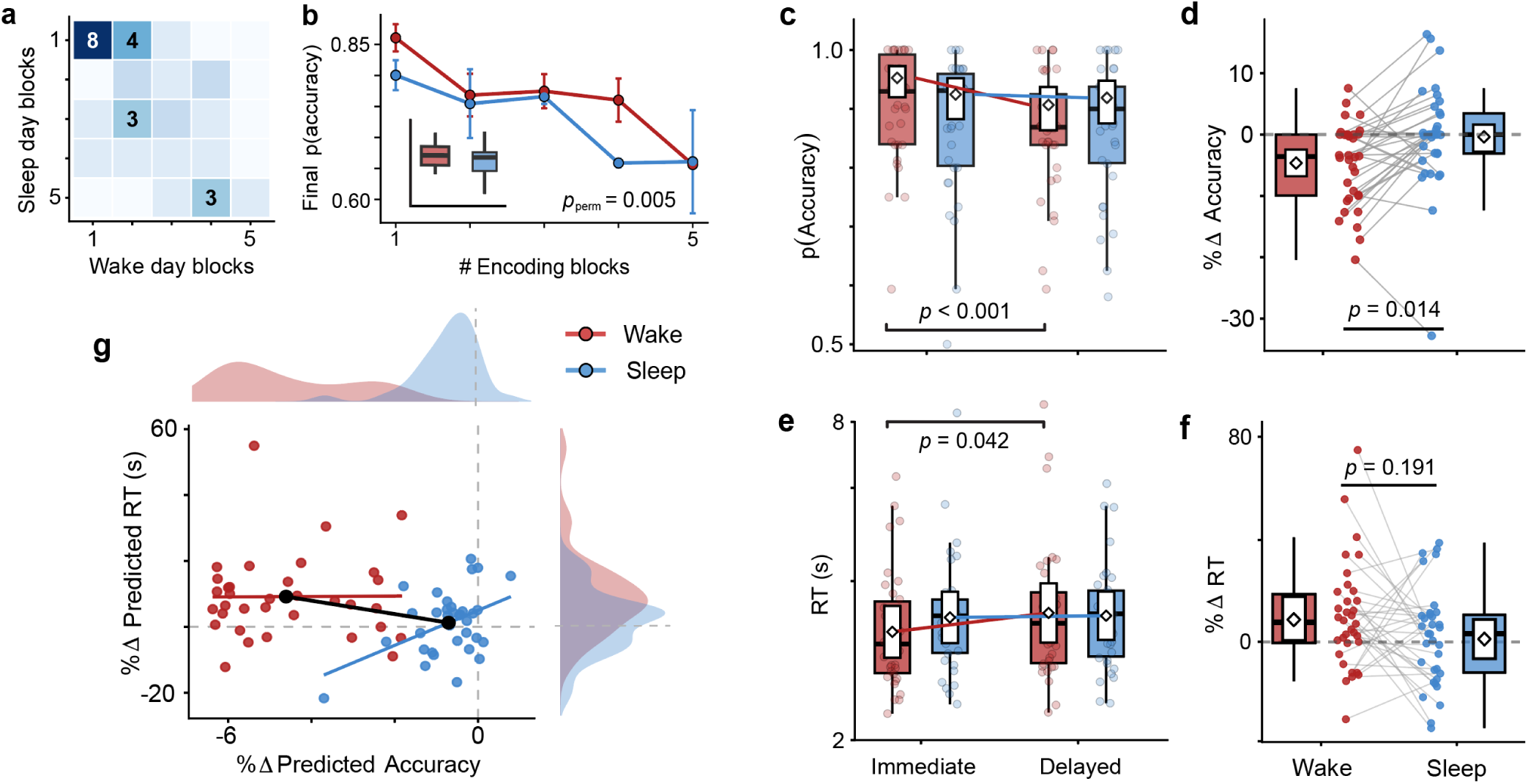
Overnight sleep preserves newly learned declarative memory. **a**, Joint distribution of the number of encoding blocks completed on Wake and Sleep days (*n* = 34). Cell shading indicates the number of participants. **b**, Final-block encoding accuracy as a function of the number of encoding blocks completed on Wake and Sleep days (mean ± s.e.m.; *P*_perm_ = 0.005). Inset, paired comparison of final-block encoding accuracy between conditions (*P*_perm_ = 0.019). **c**, Recall accuracy at immediate and delayed retrieval on Wake and Sleep days. Points and boxplots show participant-level values; white diamonds and boxes show binomial-GLMM-estimated marginal means and 95% confidence intervals (CIs), respectively. Accuracy declined across wakefulness (*P <* 0.001) but not across sleep (*P* = 0.577). **d**, Relative delayed-to-immediate change in recall accuracy for Wake and Sleep days. Colored points show participant-level changes; white diamonds and boxes show binomial-GLMM-derived relative changes and 95% CIs. The Condition × Test phase interaction was significant (*P* = 0.014). **e,f**, Retrieval latency (**e**) and relative delayed-to-immediate change in retrieval latency (**f**), shown as in **c,d**using a Gamma GLMM. Retrieval latency increased across wakefulness (*P* = 0.042) but not across sleep (*P* = 0.831); the Condition × Test phase interaction was not significant (*P* = 0.191). **g**, Relationship between participant-specific changes in recall accuracy and retrieval latency derived from the fitted GLMMs. Points show response-scale relative changes, with marginal distributions shown above and at right; black points indicate condition means. Accuracy and retrieval-latency changes were not significantly associated in either condition (both *P*_perm_ *>* 0.05). Within-condition contrasts in **c,e** used Wald *z* tests of GLMM-estimated marginal contrasts; interactions in **d,f** used likelihood-ratio tests of nested GLMMs.

Retrieval latency showed a similar temporal pattern: responses slowed from immediate to delayed testing after wakefulness but remained stable following sleep (Fig. 2e), although the corresponding difference in change between conditions was not significant (Fig. 2f). Changes in recall accuracy and retrieval latency were unrelated within either condition (Fig. 2g), suggesting that memory retention and retrieval efficiency may capture distinct aspects of long-term memory performance. These findings show that sleep preserved newly learned declarative memory across the retention interval.

### Learning induces opposing aperiodic shifts in wake and sleep

This selective preservation of declarative memory across sleep raised a corresponding neural question: how does the cortical state shaped by learning during wakefulness reorganize as the brain enters sleep? Consistent with previous studies^16,32,47,49,50^, aperiodic slopes were steeper during NREM than Wake across conditions. Beyond this expected state difference, learning shifted the aperiodic 1/f slope in opposite directions across states: relative to Sham, slopes were flatter during pre-sleep Wake but steeper during early N2–N3 (Fig. 3a,b). Regional and sensor-space analyses showed that this reversal was expressed broadly across the scalp, with widespread Wake flattening followed by broadly distributed NREM steepening that was most pronounced over frontocentral cortex (Fig. 3c–h). The principal regional effects were preserved in every leave-one-participant-out (LOPO) analysis. By post-sleep Wake, no reliable Learning–Sham difference remained (Fig. 3i). Therefore, learning did not leave a persistent aperiodic displacement across brain states. Instead, the aperiodic operating regime shifted in the opposing direction as the brain entered NREM sleep. This pattern was preserved across alternative reference schemes and direct log–log fitting, and no corresponding learning-related aperiodic modulation was observed during REM sleep (Figs. S6.1–S6.2, S7.1 and S11.1).

**Figure 3:**
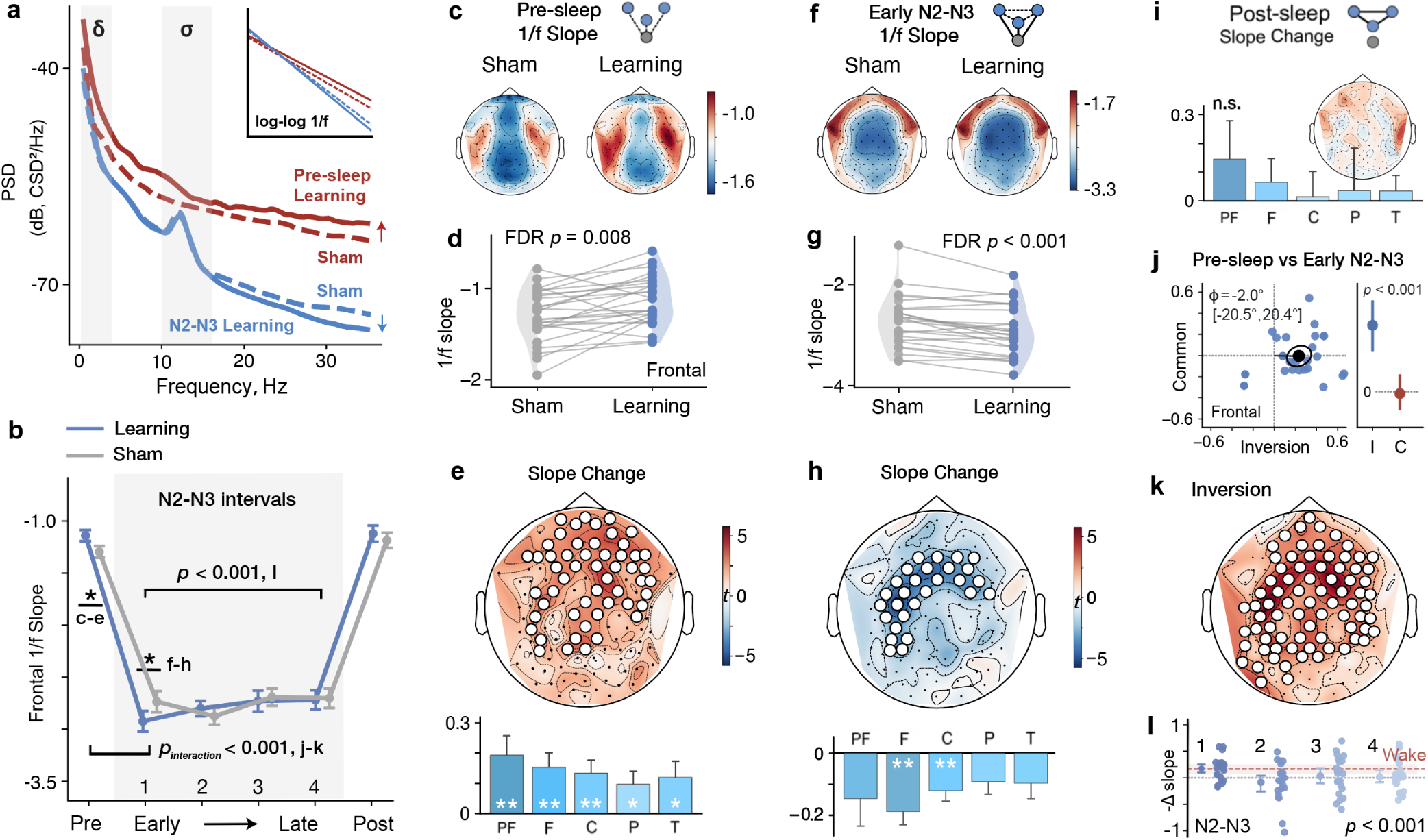
Learning induces a transient wake-to-NREM inversion of cortical aperiodic dynamics. **a**, Representative power spectra during pre-sleep Wake and early N2–N3 under Learning and Sham conditions. Inset shows the corresponding aperiodic fits on a log–log scale. **b**, Temporal trajectory of frontal region of interest (ROI) aperiodic slope across pre-sleep Wake (n = 26), four 30-min N2–N3 intervals (n = 25) and post-sleep Wake (n = 6). The NREM intervals correspond to the first two and final two available 30-min intervals within each overnight recording. Learning-related slope changes reversed from pre-sleep Wake to the first NREM interval (*n* = 23*, P*_perm_ *<* 0.001), and the early-NREM effect was stronger than the mean effect across the three later intervals (*n* = 25*, P*_perm_ *<* 0.001). Lines and error bars show condition means and s.e.m. **c–e**, Pre-sleep Wake aperiodic slopes (n = 26). **c**, Condition-mean topographies. **d**, Participant-level frontal-ROI slopes, showing Learning-related flattening (*q*_FDR_ = 0.008). **e**, Learning-minus-Sham effects across electrodes, displayed as a one-sample *t*-statistic map (top) and five predefined ROIs (bottom; mean ± s.e.m. across participants). Flattening was significant across all five ROIs (*q*_FDR_ ≤ 0.033) and formed a significant sensor-space cluster (*P*_cluster_ = 0.002). **f–h**, Corresponding analyses during early N2–N3 (n = 25). Learning produced steepening in the frontal and central ROIs (*q*_FDR_ *<* 0.001 and *q*_FDR_ = 0.006, respectively) and a significant frontocentral cluster (*P*_cluster_ = 0.002). **i**, Post-sleep Wake Learning-minus-Sham effects (n = 6). No ROI or cluster reached significance. **j**, Orthogonal decomposition of paired frontal-ROI Wake and early-NREM effects into an inversion mode, capturing opposite-direction changes across states, and a common mode, capturing same-direction changes (*n* = 23). Blue points show participant-level coordinates, the black point the observed group centroid and ellipses the participant-bootstrap distribution of the centroid. The group vector was closely aligned with the flatter-Wake, steeper-NREM inversion axis (*φ* = 2.0°, 95% bias-corrected and accelerated (BCa) CI, 20.5°–20.4°), with a significant positive projection onto this axis (*P*_perm_ *<* 0.001). **k**, Electrode-wise Wake-minus-NREM contrast showing a widespread positive inversion effect (*P*_cluster_ *<* 0.001). **l**, Temporal expression of the NREM counter-shift across intervals after sign reversal of the NREM effects. The counter-shift was strongest during the first NREM interval and was reduced across the three later intervals (*n* = 23*, P*_perm_ *<* 0.001). Summary markers show means and 95% BCa CIs; the red dashed line and band show the pre-sleep Wake mean and 95% BCa CI. ROI effects were tested by two-sided sign-flip tests with FDR correction; sensor-space effects were tested by two-sided cluster-based permutation tests. White circles indicate electrodes belonging to significant clusters; asterisks denote FDR-significant ROIs (\**q*_FDR_ *<* 0.05; \*\**q*_FDR_ *<* 0.01). PF, prefrontal; F, frontal; C, central; P, parietal; T, temporal.

The opposing Wake and NREM effects could still reflect two separate state-specific responses rather than a state-opponent pattern expressed within individuals. To distinguish these possibilities, we decomposed each participant’s paired Wake and early-NREM effects into orthogonal inversion and common modes, separating opposite-direction from same-direction changes across states (Fig. 3j). The paired effects were preferentially expressed along the inversion dimension: the group vector lay almost directly along the flatter-Wake, steeper-NREM axis, with a significant positive projection onto this direction. The minimal common-mode displacement further indicated comparable, opposing contributions from both states rather than being dominated by a change in either Wake or NREM alone. Inversion magnitude also exceeded common-mode magnitude across participants (*P*_perm_ = 0.003): 17 of 23 participants were inversion-dominant, and 16 of these expressed the flatter-Wake, steeper-NREM direction, showing that inversion was expressed within individuals rather than emerging from different participants contributing the two state-specific effects. An electrode-wise Wake–NREM contrast showed that this state-opponent organization extended broadly across the scalp (Fig. 3k). This reversal was temporally concentrated near sleep onset, with the opposing sleep effect strongest early in NREM and diminishing as sleep progressed (Fig. 3l). Thus, the observed two-state pattern was expressed within participants as a distributed cross-state inversion, consistent with a rapid, experience-dependent reorganization of cortical activity upon entry into NREM sleep.

### Aperiodic shifts dissociate from oscillations

The robust opposing aperiodic shifts across Wake and NREM raised a key question about conventional band-limited readouts. Wake theta and NREM SWA and sigma are widely used markers of learningand sleep-related physiology, but their power contains both narrowband oscillatory and aperiodic contributions. Could learning-related changes in these bands reflect the broadband 1/f shift, or persist after removal of the aperiodic component? Once this component was removed, Learning and Sham spectra showed little separation across Wake theta and NREM SWA and sigma bands (Fig. 4a,b). Consistent with this, none of the three residual-power measures showed reliable learning-related modulation across predefined ROIs or sensorspace clusters (Fig. 4c,e,f,h,i,k). Thus, the opposing aperiodic shifts across Wake and NREM were not accompanied by parallel changes in theta, SWA or sigma power after accounting for the 1/f background.

**Figure 4:**
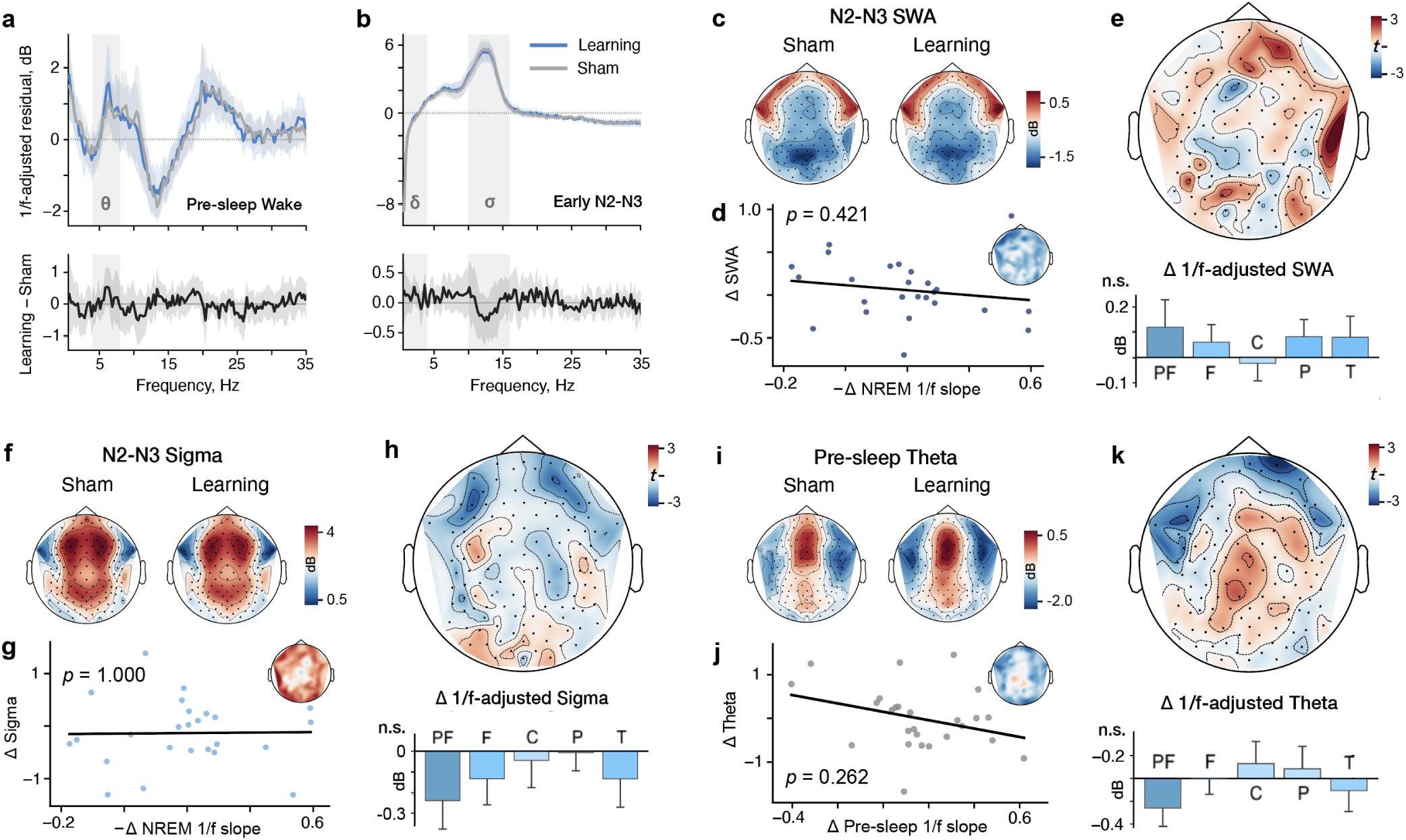
Learning-related aperiodic shifts dissociate from oscillatory power. **a,b**, Aperiodic-adjusted residual spectra (dB) during pre-sleep Wake (**a**, *n* = 26) and early N2–N3 (**b**, *n* = 25). Upper panels show participant-equal Learning and Sham means; lower panels show paired Learning-minus-Sham differences. Shading indicates 95% Student-*t* CIs across participants. Gray bands denote Wake theta (4–8 Hz), NREM SWA (0.5–4 Hz) and NREM sigma (10–16 Hz); spectra were binned at 0.25 Hz for display. **c–e**, Early N2–N3 aperiodic-adjusted SWA power. **c**, Condition-mean topographies. **d**, Across participants, frontal-ROI NREM aperiodic and SWA effects were not reliably associated (*P*_bonf_
= 0.421). Inset shows electrode-wise Spearman correlations across participants, displayed from -1 to 1. **e**, Learning-minus-Sham effects across electrodes, displayed as a one-sample *t*-statistic map (top) and five predefined ROIs (bottom). **f–k**, Corresponding analyses for early N2–N3 sigma power (**f–h**) and pre-sleep Wake theta power (**i–k**), shown as in **c–e**. For **e,h,k**, ROI Learning-minus-Sham effects were tested by two-sided participant sign-flip tests with FDR correction across the five predefined ROIs; none survived correction (SWA, all *q*_FDR_ ≥ 0.486; sigma, all *q*_FDR_ ≥ 0.457; theta, all *q*_FDR_ ≥ 0.508). Sensor-space effects were tested by two-sided cluster-based permutation tests; no cluster was identified or reached significance. For **d,g,j**, Spearman correlations were tested using two-sided permutation tests and Bonferroni-corrected across the three comparisons.

However, even weak group-level oscillatory changes could covary with the aperiodic effect across individuals, in which case individual differences in aperiodic reorganization might simply track conventional oscillatory physiology. This was not supported: participants showing larger NREM aperiodic changes did not systematically show larger SWA or sigma changes, and Wake aperiodic changes were similarly unrelated to theta-power changes (Fig. 4d,g,j). Electrode-wise correlation maps likewise showed no coherent spatial correspondence between the aperiodic and oscillatory learning effects. Together, these findings indicate that the learning-related reorganization of aperiodic dynamics was not mirrored by corresponding oscillatory changes, either at the group level or across individuals.

### Learning reshapes cortical population geometry

At the population level, the reciprocal Wake–NREM aperiodic response could take different forms. Learning might simply shift the central tendency of aperiodic activity across the cortex, or it might reorganize the broader operating regime in which cortical activity was expressed. Participant-level median slopes established the direction of the effect but did not reveal how moment-to-moment cortical states are distributed around that shift. We first examined the distribution of aperiodic slopes. Learning produced a broad displacement toward flatter slopes during pre-sleep Wake and toward steeper slopes during early N2–N3 (Fig. 5a,b), indicating that learning shifted the distribution of aperiodic states occupied during Wake and NREM.

**Figure 5:**
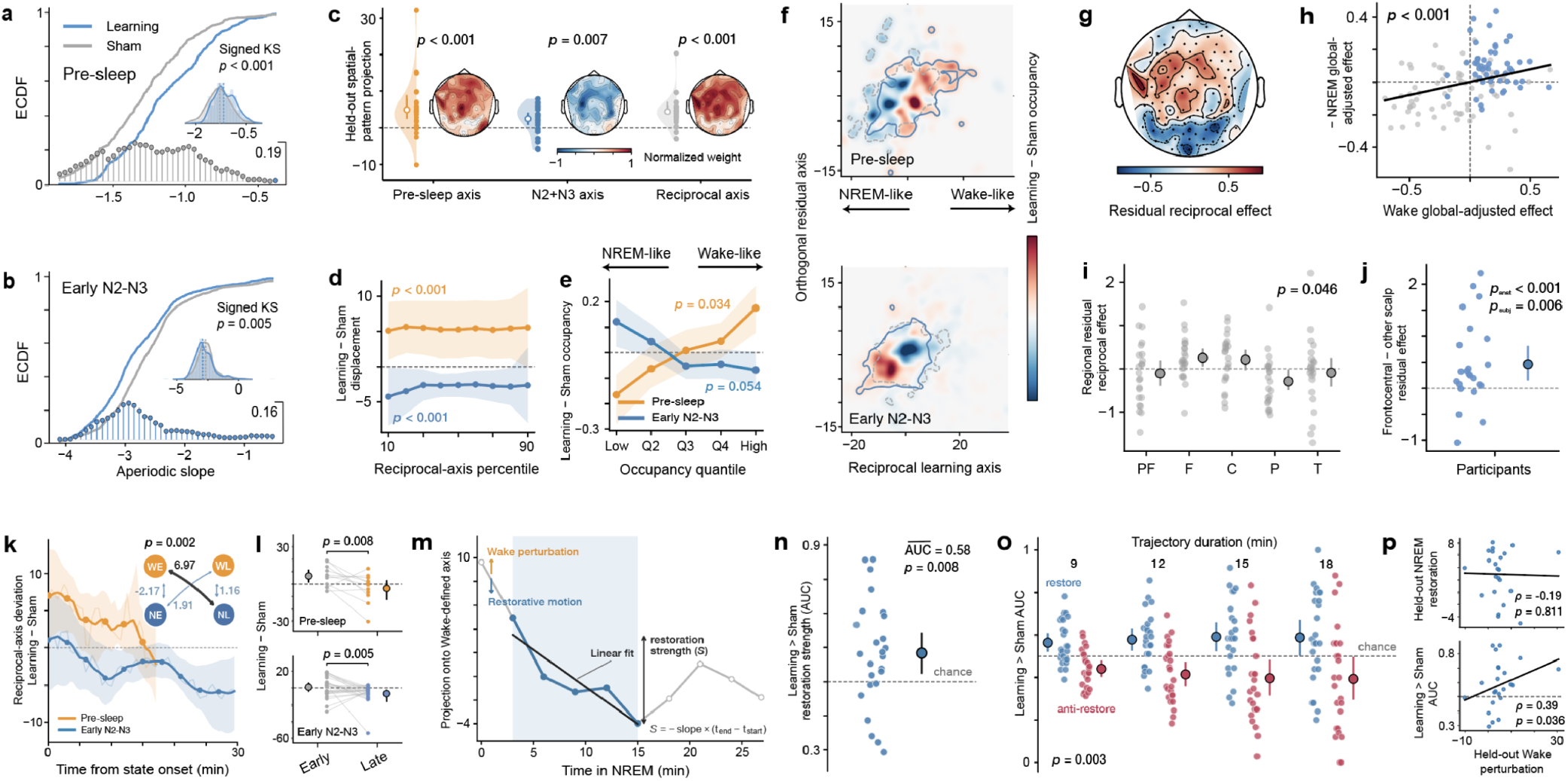
Cross-state population geometry reveals structured renormalization of cortical operating regimes. **a,b,** Participant-equal empirical cumulative distribution functions (ECDFs) of frontal-ROI aperiodic slope during pre-sleep Wake (**a**, *n* = 26) and early N2–N3 (**b**, *n* = 25). Curves show participant-equal ECDFs; lollipops show pointwise Learning-minus-Sham differences (Δ ECDF; right y-axis), and insets show slope distributions. Learning shifted slopes toward flatter values during pre-sleep Wake (*P*_perm_ *<* 0.001) and steeper values during early N2–N3 (*P*_perm_ = 0.005; participant-permutation tests of the signed Kolmogorov–Smirnov statistic). Full distributions across electrodes are reported in Fig. S9.1. **c,**Leave-oneparticipant-out (LOPO) generalization of pre-sleep, N2–N3 and reciprocal population axes. Topographies show normalized mean axis weights. Pre-sleep (*n* = 26), N2–N3 (*n* = 25) and reciprocal (*n* = 23) patterns generalized out of sample (all *P*_perm_ *<* 0.01, full-refit participant permutation tests). **d,** Learningminus-Sham displacement across the 10th–90th percentiles of the held-out reciprocal-axis distribution. Positive values indicate Wake-like and negative values NREM-like displacement. Displacement differed from zero during pre-sleep Wake (*P*_perm_ *<* 0.001) and early N2–N3 (*P*_perm_ *<* 0.001). **e,** Learning-minus-Sham occupancy differences across five equal-probability bins of the held-out reciprocal axis; whole-profile permutation omnibus: pre-sleep, *P*_perm_ = 0.034; early N2–N3, *P*_perm_ = 0.054. **f,** Participant-equal Learning-minus-Sham occupancy differences in a common two-dimensional population-state space. Color denotes occupancy difference and contours show condition densities. **g,** Residual reciprocal topography after removal of each participant’s whole-scalp mean. **h,**Global-mean-adjusted Wake and sign-reversed NREM effects were spatially correlated across 121 electrodes (*ρ* = 0.304, *P*_perm_ *<* 0.001; electrode-label permutation test). **i,** Residual reciprocal effects differed across five predefined ROIs (*P*_perm_ = 0.046; joint participant sign-flip omnibus). PF, prefrontal; F, frontal; C, central; P, parietal; T, temporal. **j,** The residual reciprocal effect was stronger over frontocentral than other electrodes across participants (*P*_participant_ = 0.006; one-sided participant sign-flip test) and exceeded randomly reassigned electrode sets (*P*_anatomical_ *<* 0.001; one-sided). **k,** Time-resolved Learning-minus-Sham reciprocal-axis deviation during the first 30 min of Wake and early N2–N3. Inset shows early/late cross-state correspondence; early-Wake/late-NREM correspondence exceeded the other three pairings (*P*_perm_ = 0.002). WE/WL, early/late Wake; NE/NL, early/late NREM. **l,** Learningminus-Sham reciprocal-axis deviation decreased from early to late during pre-sleep Wake (*P*_perm_ = 0.008, *n* = 11 with ≥ 10 min of Wake available) and early N2–N3 (*P*_perm_ = 0.005, *n* = 25). **m,** Label-blind restoration analysis of NREM trajectories projected onto a Wake-defined axis; restorative sequences required a negative fitted slope and positive net drop. **n,** Area under the curve (AUC) quantifies the probability that restoration strength was greater in Learning than Sham; mean AUC = 0.583 (95% BCa CI 0.521, 0.642, *P*_perm_ = 0.008; one-sided). **o,**Direction specificity across trajectory durations. Learning preferentially increased Wake-opposing rather than Wake-directed motion (restore-minus-anti AUC, *P*_omnibus_= 0.003; one-sided). **p,** Relationship of heldout Wake perturbation magnitude to subsequent NREM reorganization. Top, static opposing NREM displacement; bottom, Learning-specific restorative trajectory expression quantified by restoration AUC. Larger Wake perturbation was not associated with static opposing NREM displacement (*ρ* = − 0.19, *P*_perm_ = 0.811) but was associated with greater restoration AUC (*ρ* = 0.39, *P*_perm_ = 0.036); one-sided Spearman permutation tests. Points show participants or held-out participant values as applicable; summary markers and shading show means and 95% participant-bootstrap BCa CIs, except in **d**, where shading shows a simultaneous 95% bootstrap confidence band across percentiles. Two-sided full-refit participant permutation tests were used in **c,d,k,l**.

We therefore asked whether this distribution-wide shift was structured along a reproducible cortical population geometry. Multichannel Learning-minus-Sham patterns defined from other participants generalized to held-out individuals in both Wake and NREM. More importantly, a reciprocal axis defined by the opposing Wake and NREM effects also generalized out of sample (Fig. 5c). This spatial organization was robust across split epoch halves and participants (Fig. S10.1). Thus, the cross-state reversal represented a shared cortical population dimension, rather than a pattern that emerged only after averaging across participants. Along this reciprocal dimension, Learning displaced population activity across much of the underlying distribution, toward the Wake-like direction before sleep and toward the opposing NREM-like direction after sleep onset (Fig. 5d). This was accompanied by a redistribution in the occupancy of reciprocal population states (Fig. 5e,f). Our findings indicate that learning did not simply change the magnitude of aperiodic activity, but displaced the cortical population operating regime, with subsequent NREM shifting that regime in the opposing direction.

Beyond this global reciprocal shift, the Wake–NREM reorganization retained a reproducible spatial structure. To test this, we removed each participant’s mean whole-scalp effect and examined the residual reciprocal pattern. The opposing Wake and NREM effects remained spatially structured after this adjustment, and electrodes showing stronger learning-related perturbation during Wake preferentially expressed a stronger response in the opposing direction during NREM, revealing a spatially selective component of cross-state renormalization (Fig. 5g,h). This spatial selectivity was organized regionally, with the residual reciprocal effect differing across predefined cortical regions and showing the strongest expression over frontocentral cortex (Fig. 5i,j). This frontocentral predominance reflected contributions from both learning-related Wake flattening and opposing NREM steepening (conjunction *P* = 0.030), was reproducible across participants, and exceeded that expected from randomly reassigned electrodes. Thus, cross-state renormalization was not a uniform whole-cortex reversal, but a spatially structured population response.

### Wake perturbations organize restorative sleep dynamics

We next asked whether the opposing Wake and NREM effects were temporally coordinated, rather than reflecting two static consequences of learning. The learning-induced Wake displacement was strongest immediately after learning and subsided toward sleep, whereas opposing NREM expression strengthened after sleep onset (Fig. 5k,l). The spatial organization followed the same temporal sequence: the pattern expressed early in Wake corresponded most strongly to the opposing pattern that emerged later in NREM, exceeding the three alternative cross-state temporal pairings (Fig. 5k). The early-to-late relationship remained robust across temporal-edge definitions (10–50%), showing that the reciprocal population organization continued to evolve over time beyond the initial state-transition inversion.

These early-versus-late analyses nevertheless reduced the continuous trajectory to two temporal windows and could not establish whether NREM dynamics were specifically linked to the preceding Wake perturbation. We therefore asked whether NREM dynamics were specifically organized with respect to that perturbation. NREM trajectories were projected onto a held-out, Wake-defined perturbation axis with NREM condition labels concealed, and sustained movement opposing this prior-Wake direction was quantified as restoration strength (Fig. 5m). After condition labels were revealed, restoration was selectively stronger following Learning than Sham and discriminated between the two conditions (mean within-participant AUC = 0.583, Fig. 5n). This was not simply because Learning produced greater NREM movement irrespective of direction: the additional movement was preferentially directed opposite to the prior Wake perturbation rather than toward it, and this directional bias was robust to trajectory length (Fig. 5o). An exact test across all possible Wake condition-label assignments showed that the true learning-defined Wake geometry produced stronger Learning-specific expression of restorative NREM trajectories than randomized source geometries (*P* = 0.045). This direction-specific organization also persisted when Learning and Sham trajectories were matched for their initial position along the Wake-defined axis and for NREM time (mean difference = 2.400, 95% BCa CI 0.387–4.250, *P* = 0.011), indicating that NREM trajectories were specifically organized with respect to the preceding Wake geometry rather than reflecting generic movement during sleep, regression to the mean, or current position alone.

Importantly, coupling to prior Wake experience was expressed in the temporal organization of NREM dynamics rather than as a simple scaling of two state-level effects. The size of the held-out Wake perturbation did not predict the overall opposing NREM displacement. Instead, larger Wake perturbations predicted stronger Learning-specific restorative trajectories directed against the preceding perturbation (Fig. 5p). Across spatial, temporal and trajectory-level analyses, cross-state renormalization emerged as a temporally evolving response that was systematically organized with respect to the preceding learning-related Wake perturbation, consistent with a restorative process during NREM sleep.

### Wake–sleep renormalization predicts consolidation

So far, we have shown that sleep selectively preserves declarative memory relative to wakefulness (Fig. 2), and that learning induces a transient Wake-to-NREM inversion of cortical aperiodic dynamics (Fig. 3). This reciprocal reorganization was expressed most robustly in the aperiodic component (Fig. 4) and extended to a temporally evolving, spatially structured population pattern that generalized to held-out participants (Fig. 5). We finally asked whether individual differences in this wake-to-NREM reorganization tracked the benefits of sleep-dependent memory consolidation. Greater learning-related flattening of the aperiodic slope during pre-sleep Wake was associated with greater sleep-specific memory benefit, whereas the corresponding NREM relationship was weaker and opposite in direction (Fig. 6a). Given the reciprocal organization established above, we next considered the two states jointly. Greater expression of the static Wake–NREM inversion was associated with greater sleep-specific memory benefit (Fig. 6b,c). As a state-specificity control, REM slope differences were unrelated to sleep-specific memory benefit (Fig. S12.1).

**Figure 6:**
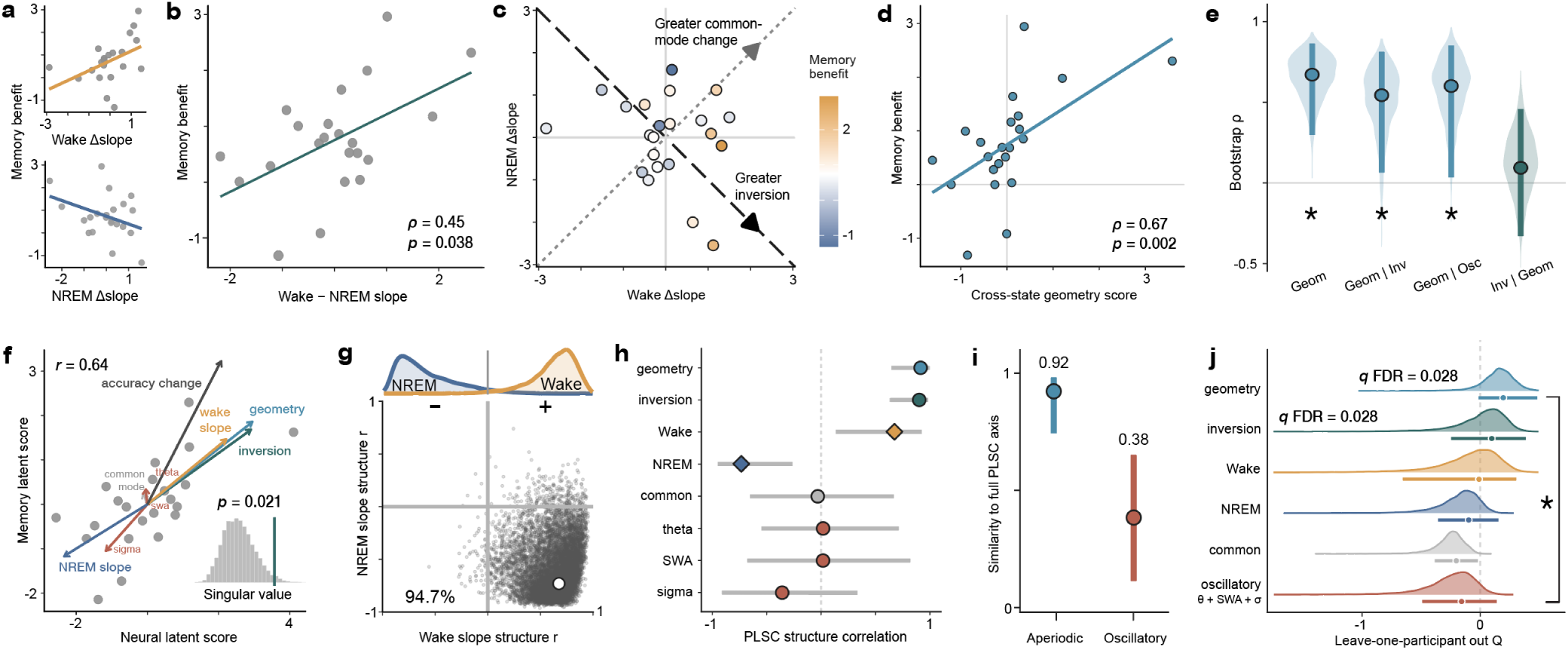
Cross-state renormalization of aperiodic operating regimes predicts memory consolidation. **a,** Sleep-specific memory benefit, defined as the sleep–wake difference in delayed-minus-immediate change in empirical-logit recall accuracy, and its relationship to Learning-minus-Sham aperiodic slope changes during pre-sleep Wake and early NREM sleep (Wake: *ρ* = 0.474, *P*_perm_ = 0.028; NREM: *ρ* = − 0.202, *P*_perm_ = 0.358). **b,** Sleep-specific memory benefit was positively associated with static Wake–NREM inversion, the orthogonal Wake-minus-NREM contrast constructed from standardized Wake and NREM slope composites (*ρ* = 0.452, *P*_perm_ = 0.038). **c,** Joint wake–NREM geometry of the state-specific aperiodic slope changes. The dashed diagonal denotes the inversion axis and the dotted diagonal the orthogonal common-mode axis. **d,** Sleep-specific memory benefit was positively associated with the cross-state geometry score, a LOPO projection of each participant’s Learning-related Wake-to-NREM population displacement onto the reciprocal spatial axis defined from the remaining participants (*ρ* = 0.671, *P*_perm_ = 0.002). **e,** Participant-bootstrap distributions of the geometry score–memory benefit association. The association remained after adjustment for static inversion (partial *ρ* = 0.559, 95% BCa CI, 0.064–0.811; *P*_perm_ = 0.010) and for pre-sleep theta, NREM SWA and NREM sigma (partial *ρ* = 0.601, 95% BCa CI, 0.026–0.849; *P*_perm_ = 0.008), whereas static inversion showed little residual association after adjustment for geometry score (partial *ρ* = 0.068, 95% BCa CI, 0.329 to 0.421; *P*_perm_ = 0.773). **f,** Partial least squares correlation (PLSC) analysis of the multivariate neural relationship with sleep-specific memory benefit. Neural and memory latent scores were correlated (*r* = 0.644); the latent mode was significant by singular-value permutation (singular value = 0.885, *P*_perm_ = 0.021). The neural latent axis was stable to LOPO refitting (mean cosine similarity = 0.993, range = 0.957–1.000; all folds *>* 0.90). Arrows show correlations of neural features and behavioral outcome with the latent scores; Wake and NREM aperiodic slope projections were not included as fitted predictors and are shown as supplementary features. **g,** Participant-bootstrap geometry of the Wake and NREM slope projections onto the PLSC latent mode; gray points show bootstrap estimates and the white point the observed full-sample projection; 94.7% of samples showed opposite signs. **h,** PLSC structure correlations. **i,** Feature-family preservation of the full PLSC neural axis, quantified as cosine similarity after refitting with either aperiodic features (geometry score, static inversion and common mode) or oscillatory features (theta, SWA and sigma). **j,** LOPO predictive performance for neural predictors, quantified by held-out *Q*^2^. Ridges show participant-bootstrap distributions. Geometry score and static inversion survived FDR correction (both *q*_FDR_ = 0.028). In **e,h–j**, summary markers indicate observed estimates and 95% BCa CIs. All analyses included *n* = 22 participants.

A scalar Wake–NREM contrast, however, does not retain the spatial organization of the underlying population shift. Building on the reciprocal model established in Fig. 5, we quantified each participant’s expression of this spatial pattern as the cross-state geometry score, a behavior-independent measure. For each participant, the reciprocal Wake-to-NREM axis was defined exclusively from the neural data of the remaining participants, and the held-out participant was then projected onto this independently defined axis. The geometry score was strongly related to sleep-specific memory benefit (Fig. 6d), and this association remained positive and significant in every LOPO reanalysis (*ρ* = 0.625–0.742; all *P*_perm_ ≤ 0.003). The relationship also persisted after accounting for both the simpler static inversion and oscillatory measures, whereas static inversion showed little residual relationship with memory once geometry was taken into account (Fig. 6e). Our findings indicate that sleep-specific memory benefit was captured not only by the magnitude of cross-state inversion, but by its reproducible multichannel organization across the cortical population.

We next evaluated whether sleep-dependent memory benefit was organized around a broader multivariate neural pattern rather than any single predefined measure. Partial least squares correlation (PLSC) identified a significant latent mode linking neural variation to memory outcome (Fig. 6f), and this neural axis was highly reproducible under LOPO refitting. Supplementary Wake and NREM aperiodic-slope projections aligned in opposite directions with the memory-related latent axis (Fig. 6g), indicating that greater memory benefit was associated with a reciprocal organization of Wake and NREM aperiodic dynamics, which was not specified a priori by the multivariate model but emerged from the joint neural–behavioral covariance structure. The structure of the mode revealed a coordinated cross-state organization: geometry and static inversion were the dominant contributors to the latent axis, whereas common-mode and oscillatory measures contributed comparatively little (Fig. 6h). Refitting the PLSC using only aperiodic measures closely preserved the full neural axis, whereas an oscillatory-only model showed much weaker alignment (Fig. 6i). Thus, the memory-related signal was embedded in a multivariate cross-state aperiodic organization, with reciprocal Wake–NREM reorganization forming the principal neural dimension.

To establish whether these neural signatures generalize beyond the participants from which they were estimated, we evaluated each neural predictor using LOPO prediction of sleep-specific memory benefit (Fig. 6j). Cross-state geometry showed positive held-out predictive performance and exceeded its permutation null. Static inversion also exceeded its permutation null, whereas Wake, NREM and common-mode measures did not. The oscillatory feature family likewise showed no positive held-out performance and was significantly outperformed by cross-state geometry. These results extend the association between cross-state renormalization and memory beyond the observed sample: the neural signatures that generalized were those capturing the reciprocal relationship between wakefulness and NREM sleep, rather than either state considered in isolation. The Wake-to-NREM inversion therefore links the learning-induced displacement of waking cortical dynamics to their subsequent reciprocal expression during sleep, providing a population-level physiological axis along which the effectiveness of declarative memory consolidation varies across individuals.

## Discussion

By tracking aperiodic dynamics continuously from wakefulness into sleep, we provide evidence for perturbation-dependent renormalization as a systems-level process. Learning shifted the cortical operating regime during wakefulness, and both the temporal trajectory and cortical distribution of the subsequent opposing NREM response were organized by the preceding waking perturbation, indicating a structured restorative process spanning wakefulness and sleep. This cross-state organization generalized across individuals, and its strength predicted greater sleep-specific memory benefit in held-out participants. These observations suggest that consolidation depends not only on physiology expressed within sleep, but also on how post-learning waking dynamics are transformed during subsequent sleep. Together, they support a role for sleep in consolidation that extends beyond reducing interference^51^ to counterbalancing the population-level changes induced by learning.

Across scales, neural regulation depends on recent activity and plasticity. Excitatory synapses undergo homeostatic downscaling during sleep^9–11^, whereas neuronal firing rates recover toward cell-specific set points after sustained sensory perturbation^18,52^. At larger scales, learning locally increases subsequent SWA over task-engaged cortex^21^, and waking experience displaces network dynamics from criticality before sleep restores them^19^. For a plastic brain, this creates a broader control problem: once experience has changed the system, subsequent regulation must proceed in relation to what changed. Our findings place this problem at the level of population dynamics across wakefulness and sleep. Across space, regions with larger waking perturbations showed stronger opposing NREM responses rather than a uniform whole-brain reset. Across time, NREM dynamics evolved along a population axis defined during wakefulness. Across individuals, larger waking perturbations predicted stronger restorative trajectories. More importantly, restoration was not determined by current state alone. Even after matching Learning and Sham trajectories for position along the Wake-defined axis and NREM time, Learning trajectories showed stronger motion opposing the preceding waking perturbation. Thus, subsequent NREM dynamics depended not only on where the system was, but on how it had arrived there, giving perturbation-dependent renormalization the logic of history-dependent negative feedback. In this framework, homeostasis is expressed not only in the recovery of individual neural variables, but in how population activity moves through neural state space after learning. The wake-to-sleep transition itself therefore becomes part of the restorative process. This raises a deeper question: why should learning and subsequent sleep push cortical dynamics in opposing directions?

The waking displacement may provide part of the answer. Learning requires a neural system to modify itself while continuing to process incoming information, which creates a tension between plastic change and dynamical stability^1,2^. The displacement observed here may reflect not a failure of regulation, but a transient consequence of learning-induced circuit reorganization. One possibility is that spectral flattening accompanies faster population dynamics with shorter temporal integration windows^53^ and a greater weighting toward excitation^42^. These conditions can alter neuronal responsiveness^54^ and accelerate cortical information processing^55^. The transient nature of the learning-induced waking displacement in our data is consistent with this possibility. None of these physiological mappings is unique to the aperiodic slope^40,41^, but together they motivate an explanation in which learning may temporarily bias cortical dynamics toward change rather than stability, making the observed perturbation a dynamical cost of modifying the system.

From this perspective, the opposing NREM response should not simply undo what learning has done. It may instead counterbalance a computational bias introduced by learning. If learning temporarily biases cortical dynamics toward rapid processing and modification, the subsequent steepening may reflect a complementary shift toward relatively more temporally correlated^53^, inhibition-weighted^40,42,56^, and lower-complexity population activity^57^. Such conditions could provide a regime favorable to the integration and stabilization of recently acquired information^58,59^. The cross-state time course supports this view: the learning-related effect evolved from maximal flattening immediately after learning to opposing steepening in early NREM, which then diminished later in the night. Importantly, the NREM trajectory varied with recent plastic history rather than scaling with the static waking displacement. Perturbation-dependent renormalization therefore reflects a continuous trajectory through neural state space whose direction is constrained by the preceding learning perturbation, rather than a fixed difference between Wake and NREM or a history-insensitive return toward baseline. Such reorganization may shift population dynamics toward conditions compatible with stabilization.

The behavioral relevance of this cross-state reorganization suggests that renormalization and consolidation are functionally coupled. Extending prior work linking REM-dependent recalibration to overnight retention and subsequent NREM physiology^39^, we found that memory benefit was better captured by the reproducible population geometry of the Wake-to-NREM transformation than by state-specific effects or a scalar cross-state reversal. These findings position cross-state transformation as an organizing principle of consolidation: perturbations established in one state may shape population reorganization in the next. Viewed in this way, synaptic renormalization, oscillatory coordination and cross-state population reorganization describe complementary levels of sleep-dependent regulation. Synaptic changes may contribute to the evolving population trajectory, while reactivation coordinated by slow oscillations, spindles and ripples^7,13,14^ operates within, and is possibly constrained by, the history-dependent population regime indexed by aperiodic activity across wakefulness and sleep.

This distinction also changes how learning-related sleep oscillations and local sleep effects are interpreted^21,22,60^. Conventional band-limited measures conflate oscillatory and aperiodic contributions^46^, but after removing the aperiodic component, theta, SWA and sigma showed no reliable learning-related modulation. Instead, the learning-induced modulation was expressed primarily in broadband population dynamics. Recent evidence that 1/f fluctuations can inflate putative awake ripple detections further underscores the importance of this analytical separation when attributing effects to specific oscillations^48^.

A broader implication is that neural adaptation depends not only on occupying favorable operating regimes, but also on regulating transitions between them as computational demands change^61–64^. Recent evidence that the wake-to-sleep transition follows an organized dynamical trajectory indicates that falling asleep is not simply a categorical change of state^65^, and our findings suggest that recent plastic history organizes its subsequent course through NREM. This organization resembles homeorhesis, in which stability is maintained by returning toward a characteristic trajectory after perturbation rather than to a fixed point^66,67^. Here, however, the regulated object is a family of history-dependent trajectories rather than a single recovery path. From this perspective, learning capacity is constrained by the ability of a neural system to renormalize its dynamics following perturbation. Sleep may function as a dynamical operator that maps recent plastic history onto restorative population trajectories, preserving learned structure while re-establishing dynamical conditions compatible with further learning. Systems that more tightly align recovery trajectories with prior perturbations should tolerate greater plastic change without loss of stability, whereas failures of this regulation should appear as dysregulated transitions^62^, including excessive learning-induced displacement or delayed counter-regulation during sleep, rather than an abnormal operating state alone. Consistent with this framework, sleep restores subsequent encoding capacity^68–70^. We predict that the same cross-state organization will track future learning capacity and that varying learning demands and their timing relative to sleep will generate distinct waking perturbation axes with corresponding sleep trajectories. More broadly, perturbation-dependent renormalization provides a candidate control principle by which adaptive neural systems reconcile plasticity with stability. Testing these predictions across repeated wake–sleep cycles will establish how broadly this principle applies.

Several considerations should guide this interpretation. Wake recordings began after active learning, preventing us from determining whether aperiodic flattening accumulated continuously during encoding, and the limited post-sleep sample precluded a well-powered test of subsequent recovery in waking dynamics. In addition, the aperiodic slope estimated from scalp EEG does not uniquely identify the cellular mechanisms underlying the observed transformation. Finally, because neither the waking perturbation nor the subsequent sleep response was experimentally manipulated, their relationship with memory does not establish causal mediation. Simultaneous measurements across synaptic, cellular, circuit and population scales in the future could determine how local plastic changes are coordinated with the broader population-level cross-state reorganization observed here, and whether perturbation-dependent renormalization reflects a common regulatory principle across levels of neural organization. Within these boundaries, our findings support a model in which sleep reconciles stability and plasticity without requiring the system to return to a prior dynamical state, preserving what learning has encoded while renewing its capacity for further change.

## Methods

### Participants

Forty-one healthy young adults enrolled and completed the study protocol after providing written informed consent. Of these, 36 contributed valid data to at least one behavioral or neural analysis (aged 18–25 years; *M* = 23.37, *SD* = 1.99; 20 female). Participants were recruited through physical flyers, email correspondence, and the UMass SONA system, and received either monetary compensation or course credit. All participants were right-handed, fluent English speakers with normal or correctedto-normal vision, and reported no history of neurological or psychiatric disorders or use of medication.

Behavioral analyses included 34 participants. Neural analyses used analysis-specific complete samples: 26 participants for pre-sleep wake analyses, 25 for NREM analyses, and 6 for post-sleep wake analyses. Exclusion criteria included insufficient state-specific EEG data, unusable or irregular PSG recordings, loss of the ground electrode, and unsuccessful parameterization. Multivariate neural–behavioral analyses were restricted to the 22 participants with complete behavioral, pre-sleep wake EEG, and NREM EEG data. Detailed analysis-specific sample sizes and exclusion details are provided in Table S1.2.

### Procedure

All procedures were approved by the Institutional Review Board at the University of Massachusetts, Amherst (Protocol No. 5846). The study took place over the course of 21 days. The three conditions were scheduled on approximately days 7, 14, and 21. On day 1, participants completed a demographic survey, the Insomnia Severity Index (ISI)^71^, the Morningness-Eveningness Questionnaire (MEQ)^72^, and the Pittsburgh Sleep Quality Index (PSQI^73^; see Table S1.1). At each experimental session, participants completed the Stanford Sleepiness Scale (SSS)^74^ prior to performing the task. Each morning, participants received a daily questionnaire that asked when they went to bed the previous night and when they woke up in the morning. Demographic and questionnaire data are presented in Table S1.1. Participants were healthy and did not indicate any sleep disorders.

Experimental conditions took place in the Sleep Lab. Two conditions were overnight conditions for which participants came to the lab roughly 1 hour before their habitual bedtime. Participants completed encoding and immediate recall of the learning task or the sham task in the evening (~30 minutes), then were equipped with the HD-PSG cap (~45 minutes). Subsequently, participants were prepared for sleep, during which pre-sleep Wake EEG was recorded prior to lights-off, followed by overnight sleep (~8–9 hours; Fig. 1c). The duration of the pre-sleep Wake recording was determined by each participant’s sleep latency; no fixed waking interval was imposed to avoid delaying the natural wake-to-sleep transition. The next morning, the HD-PSG cap was removed, and participants completed the daily morning questionnaire and the SSS. Subsequently, participants either completed delayed recall (learning condition; ∼ 15 minutes) or were free to leave (sham condition).

The third condition was a wake condition in which participants came to the lab in the morning to complete the encoding and the immediate recall phases of the word-pair task. Afterward, they were instructed to go about their day while refraining from naps. Participants then returned to the lab in the evening, approximately 8-9 hours later, to complete the SSS and the delayed recall phase of the word-pair task. Condition order was counterbalanced across participants. Two equivalent word-pair sets were used across the sleep-learning and wake conditions and were counterbalanced across participants.

### Word Pair Task

Declarative memory was assessed using a word-pair learning task adapted from Wilson et al.^75^. Thirty-two semantically unrelated word pairs (e.g., FOX–SOUP; Fig. 1a) were selected from the South Florida Associative Norms database^76^. The task consisted of four phases: passive encoding, active encoding, immediate recall, and delayed recall (Fig. 1a-b). During passive encoding, participants viewed each word pair for 3 s, separated by a 2.5 s interstimulus interval. During active encoding, participants completed successive blocks in which all 32 cue words were presented once in random order and participants typed the corresponding target word. Feedback was provided for incorrect responses (2.5 s). Training continued in blocks until at least 20 pairs were correctly recalled within a block or until five blocks had been completed. Immediate and delayed recall tested all 32 cue words using the same procedure but without feedback.

In the sham task, participants viewed 32 pseudoword pairs matched in length to the real words used in the learning task (e.g., AZBH–TIGZ). To maintain attention, participants answered a simple yes/no question about each pair (e.g., “Was there a Y on the screen?”) by typing “Y” or “N”.

### Behavioral Modeling

We first preprocessed trials to ensure stable measurement. Reaction times (RTs) at or below 1 s were excluded, after which RTs were z-scored within each Participant × Condition combination, and trials with absolute z values greater than 2.5 were removed from both accuracy and RT analyses. Accuracy and RT were then aggregated per Participant × Condition (wake or sleep) × Task Phase (immediate or delayed recall) to generate descriptive summaries and figures. Final-block encoding accuracy was calculated separately for each Participant × Condition combination. Its association with the number of completed encoding blocks was tested using a linear mixed model with standardized block count and Condition as fixed effects and Participant as a random intercept.

For inferential analysis, we used generalized linear mixed models (GLMMs). Compared to linear mixed models or ANOVA on aggregated data, trial-level GLMMs effectively preserve within-participant variance and improve sensitivity. In addition, they correctly model binary accuracy data with a binomial likelihood and logit link, which handles ceiling effects and bounded outcomes. Binary accuracy was modeled with a binomial GLMM with a logit link including fixed effects of Condition, Task Phase, and their interaction, together with participant-specific random intercepts, random slopes for the Condition × Task Phase interaction, and a random intercept for Target. Because RTs are positively continuous and right-skewed, they were analyzed on correct trials using a Gamma GLMM with a log link and the same fixed and random effect structure.

To control for individual differences, we conducted sensitivity analyses including three covariates in both accuracy and RT control models: 1) encoding ability index, 2) SSS scores, and 3) word set. The encoding ability index was derived from a hierarchical binomial model fit to encoding trials, which included standardized block number as a covariate to capture overall learning trends and random intercepts for Participant × Condition and Target. Participant × Condition random intercepts were standardized and used as an ability measure in recall analyses. Sleepiness was quantified as the mean of morning and evening SSS scores and standardized across participants, and word set was included as a categorical covariate. Group-level comparisons were derived from model-estimated marginal means and contrasts. Model assumptions were evaluated with DHARMa residual simulations to assess uniformity, dispersion, and outliers.

For neural–behavioral analyses, recall accuracy in each Participant Condition Task Phase cell was converted to an empirical logit,

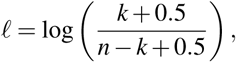

where *k* was the number of correctly recalled items and *n* was the number of tested items. Sleep-specific memory benefit for participant *i* was defined as

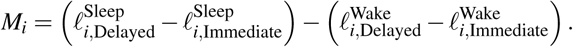

Positive values indicated greater relative preservation of recall accuracy across sleep than across wake.

### High-density Polysomnography acquisition and preprocessing

High-density PSG was acquired using a custom 128-electrode cap (Easycap, Herrsching, Germany) and BrainAmp MR plus amplifiers (Brain Products GmbH, Gilching, Germany). The EEG montage comprised 121 scalp electrodes placed according to the international 10-05 system, including an online reference at FCz, together with mastoid electrodes M1 and M2. Four EOG electrodes were placed beside and below the eyes, and 2 EMG electrodes were placed over the zygomaticus and mylohyoid muscles. Data were recorded using a hardware bandpass of DC-1000 Hz and digitized at 500 Hz using BrainVision Recorder (Brain Products GmbH, Gilching, Germany). Scalp impedances were reduced below 20 *k*Ω using high-chloride abrasive gel before the sleep intervals. All PSG analyses were conducted in Python using MNE-Python (version 1.8.0)^77^.

PSG preprocessing was performed in MNE-Python using a standardized automated pipeline. The continuous EEG signal was resampled to 200 Hz and filtered with a 60 Hz notch filter. EEG and EOG electrodes were band-pass filtered between 0.1-40 Hz, and EMG electrodes between 10-99.9 Hz. A standard 10-05 electrode montage was applied, and EEG data were re-referenced to the linked mastoids (M1 and M2) for artifact detection. Bad electrodes were identified based on the combination of variance, kurtosis, Hurst exponent, and spatial correlation metrics and were interpolated using spherical splines. On average, 6.32% of electrodes were interpolated per file, with a maximum of 26.67% of electrodes. In addition, epoch-wide and electrode-specific artifacts were automatically detected on non-overlapping 10 s segments, in which the criteria included peak-to-peak amplitude (*>* 600 *µ*V), epoch-wise and electrode-wise amplitude outliers (*z >* 5), and flat signals (*>* 2 s with SD *<* 0.2 *µ*V). If more than 10% of electrodes were contaminated within an epoch, the epoch was excluded from further analyses. Artifact segments were interpolated using spherical spline interpolation and blended with surrounding data using a Hann-tapered cross-fade window (5% of the segment length) to preserve temporal continuity.

The cleaned EEG signals and their power spectral densities were visually inspected to confirm preprocessing quality. Sleep stages were visually scored in 30-second epochs by experts according to the American Academy of Sleep Medicine criteria^78^ using the Hume toolbox in MATLAB^79^. The sleep stage characteristics were reported in Table S2.1, and the N3 proportion within the first 30 min was examined as a sleep-depth control (Fig. S2.2). After preprocessing, the EEG signals were rereferenced to the common average and transformed using a current source density (CSD) spatial Laplacian to improve spatial resolution and reduce volume conduction effects^80^. Subsequent sensor-space analyses used 121 scalp electrodes, excluding the mastoid reference electrodes M1 and M2.

### Spectral analysis

Spectral analyses were performed to quantify aperiodic slope and aperiodic-adjusted oscillatory activity during pre-sleep wakefulness, NREM sleep, REM sleep, and post-sleep wakefulness. Power spectral density (PSD) was estimated in linear power from artifact-free, non-overlapping 30-s EEG epochs using a Hann taper. To dissociate oscillatory activity from scalefree aperiodic components, PSDs were parameterized using *specparam* (version 2.0.0; formerly FOOOF)^46^, which models the power spectrum as the combination of an aperiodic 1/f component and superimposed oscillatory peaks. For all *specparam* fits, peak-width limits were 0.5-12 Hz, minimum peak height was 0, and the peak-detection threshold was 2. For the aperiodic slope analysis, six consecutive 30-s PSD estimates were averaged in linear power within each 3-minute moving window, with successive windows advanced in 30-s steps. Each window-resolved spectrum was parameterized over 1-35 Hz using a fixed aperiodic model with the knee parameter disabled. This specification was chosen because broadband fits retaining lower frequencies provide more stable exponent estimates than narrowband fits, while the fixed model provides a common single-exponent measure for comparison across wakefulness and NREM sleep and has shown better fit than the knee model during N2 and N3 sleep in scalp EEG^32^. The 1-Hz lower bound reduced sensitivity to low-frequency edge effects, resulting in a model of the form log_10_ *P*(*f*) = *b − χ* log_10_(*f*). Aperiodic slope was recorded as − *χ*, such that more negative values represented steeper spectra. For each participant, condition, electrode, and state, the median of the eligible window-resolved slope estimates was used as the summary value to reduce the influence of occasional extreme or unstable window-level fits. Learning-related effects were calculated within each participant and electrode as paired Learning-minus-Sham slope differences.

For the oscillatory analyses, the eligible 30-s PSD estimates were averaged arithmetically in linear power within each participant, condition, electrode, and state to obtain a single state-average spectrum. Averaging was performed before spectral parameterization so that each fitted spectrum represented the arithmetic mean spectral power across the selected state. Each state-average spectrum was then parameterized once over 0.5-35 Hz using specparam with a fixed aperiodic model, allowing the complete 0.5-4 Hz SWA range to be retained within the fitted spectrum. The aperiodic component obtained from this fit was used only to define the spectral background for oscillatory residual estimation and was not used as the primary aperiodic slope measure. Aperiodic-adjusted oscillatory activity was quantified from the residual spectrum, *R*(*f*) = 10 log_10_ *P*_empirical_(*f*) − log_10_ *P*_aperiodic_(*f*), expressed in decibels. Band-specific values were calculated as the arithmetic mean of residual-frequency bins within 0.5 ≤ *f <* 4 Hz for slow-wave activity (SWA), 4 ≤ *f <* 8 Hz for theta, and 10 *f <* 16 Hz for sigma. Negative residual values were retained, and residuals were not normalized by total periodic power or converted to relative power to avoid denominator coupling. Learning-related effects were calculated within each participant and electrode as paired Learning-minus-Sham differences in residual dB. For visualization of the condition-specific residual spectra in Figure 4a,b, participant-level residual spectra were aggregated into 0.25-Hz display bins before calculating group means and two-sided 95% Student-*t* confidence intervals.

Spectral model fits were visually inspected to confirm adequate parameterization. Fit quality was additionally quantified across states, conditions, and fitting ranges using *specparam* model *R*^2^ and model fit error, defined as the mean absolute difference between the empirical and full model-predicted log-power spectra across fitted frequency bins. For window-level quality control, model fit metrics were first aggregated across electrodes using the median to obtain a single fit-quality measure for each temporal window. Window-resolved fits with *R*^2^ *<* 0.80 were flagged for visual review. Twenty-seven windows (0.556% of all 4,856 windows used in the analyses) were confirmed as poor fits, and their electrode-wise slope estimates were replaced by linear interpolation from neighboring valid windows. These diagnostics are reported in Fig. S4.1. Because window-level fit metrics quantify how well the model reproduces the overall spectrum rather than whether every parameter estimate is free from isolated channel-specific artifacts or instability, electrode estimates were additionally screened using pre-specified robust distributional and spatial quality-control criteria specified in Supplementary material 5. Automatically flagged estimates were subsequently reviewed by visual inspection. Across all pre-sleep wake, NREM, and post-sleep wake spectral analyses, three of 32,186 participant-by-condition-by-electrode-by-parameter estimates (0.009%) were identified as invalid and interpolated using spherical splines separately within each participant-condition-spectral-parameter electrode map before condition contrasts were calculated. No interpolation was performed across participants or conditions. The qualitycontrol workflow and parameter-specific diagnostic panels are reported in Figs. S5.1-5.6. Sensitivity analyses omitting these window-level and electrode-level estimates produced the same conclusions.

Participants were required to contribute at least 5 min of artifact-free EEG in both the Learning and Sham conditions for state-specific wake and REM analyses. Primary pre-sleep wake analyses used up to the first 30 min of eligible post-task wake and included 26 participants for aperiodic slope and theta residual activity; analyses using the full available pre-sleep wake period were conducted as sensitivity analyses (Fig. S8.1). Primary NREM analyses used the first cumulative 30 min of artifact-free N2 and N3 epochs and included 25 participants for aperiodic slope, SWA residual activity, and sigma residual activity. This early NREM interval was selected because use-dependent changes in local sleep dynamics are largest during the initial portion of NREM sleep^21,22^. An exploratory overnight analysis characterized aperiodic slope across four cumulative 30-min N2/N3 segments, comprising the first two and final two available segments of each recording (Fig. 3b); all 25 participants contributed at least 120 min of artifact-free N2/N3 sleep. Post-sleep wake analyses were limited to aperiodic slope, used up to the first 30 min of eligible wake following morning awakening, and included six participants. The reduced sample primarily reflected recordings that ended before 5 min of usable post-awakening wake EEG had been acquired; additional exclusions resulted from loss of the ground connection or other recording failures that resulted in no usable post-sleep EEG. REM analyses used up to the first 30 min of eligible REM sleep and included participants meeting the same 5-min paired-session criterion to test whether learning-related slope changes generalized beyond NREM (Fig. S11.1).

As robustness analyses, aperiodic slope estimates were additionally tested across alternative spectral fitting frequency ranges, EEG reference schemes, and spectral fitting approaches. In addition to the primary 1-35 Hz *specparam*-derived estimates, slopes were estimated over 1-15 Hz and 15-35 Hz to assess whether the observed effects reflected broadband scalefree structure or frequency-range-specific changes. These analyses were performed using both average-reference data followed by current source density transformation and linked-mastoid reference data without CSD transformation (Fig. S6.1-6.2). To assess whether the results depended on the spectral fitting approach, we also estimated aperiodic slope using direct log-log fitting, in which a linear function was fit to the log-transformed power spectrum as a function of log frequency while omitting the 8-16 Hz range dominated by prominent oscillatory activity (Fig. S7.1). All primary results reported in the main text are based on the corresponding full-range *specparam*-derived estimates described above. Fit-quality diagnostics, REM analyses, reference and fitting-range robustness analyses, and direct log-log fitting analyses are reported in the corresponding sections of the Supplementary Material.

### Regional and sensor-space analyses

To quantify spatially specific population dynamics, we predefined five anatomically motivated regions of interest (ROIs) based on the high-density EEG layout: prefrontal, frontal, central, parietal, and temporal. The full list of electrodes included in each ROI is provided in Table S3.1 and Fig. S3.2. The frontal ROI was designated a priori as the primary representative region because the study hypotheses emphasized declarative learning-related changes in frontocentral cortical dynamics. The remaining ROIs were retained as prespecified regional comparisons, and all five ROIs were included in the multiple-comparison correction.

Participant-level electrode estimates were first obtained separately for the Learning and Sham conditions within each analyzed state. For all aperiodic and oscillatory measures, condition effects were quantified at each electrode as the paired within-participant difference, Learning minus Sham. Aperiodic effects represented differences in signed 1/f slope, whereas oscillatory effects represented differences in aperiodic-adjusted periodic power expressed in decibels. For each participant, ROI-level effects were calculated by averaging the electrode-wise condition effects across electrodes within each predefined ROI. ROI effects were tested against zero using two-sided sign-flip permutation tests of the mean with 10,000 permutations. Multiple comparisons were controlled using the Benjamini–Hochberg false discovery rate (FDR) procedure (*q* = 0.05)^81^. To assess whether significant ROI effects were driven by individual participants, we repeated the ROI analyses in a leave-oneparticipant-out analysis, excluding each participant in turn. For each iteration, two-sided sign-flip permutation tests were recomputed and the resulting permutation *P*-values were corrected across the five predefined ROIs.

Associations between aperiodic and oscillatory Learning-minus-Sham effects were assessed across participants using Spearman rank correlations. The three frontal-ROI comparisons paired NREM aperiodic slope with SWA and sigma, and pre-sleep Wake aperiodic slope with theta. Statistical significance was tested using two-sided permutation tests with Bonferroni correction. Electrode-wise correlations across participants were calculated for the corresponding spatial patterns.

For topographic inference, participant-level electrode-wise Learning-minus-Sham effect maps were analyzed using twosided one-sample cluster-based permutation tests^82^ with 10,000 participant-level sign flips. Candidate clusters were formed at an uncorrected two-sided electrode threshold of *p <* 0.05. A minimum cluster size of four electrodes was used to exclude isolated or very small electrode groupings while retaining sensitivity to spatially focal effects. The inferential conclusions were unchanged across minimum cluster-size thresholds ranging from 1 to 20 electrodes. Spatial contiguity was defined using the binary adjacency matrix derived from the scalp montage. Positive and negative clusters were formed separately, and cluster mass was defined as the sum of the signed one-sample *t*-statistics across electrodes within each cluster. Cluster-level *P* values were obtained from the permutation distribution of the maximum absolute cluster mass. Clusters with permutation *p <* 0.01 were considered significant. This stricter cluster-level threshold was adopted as a conservative criterion for the spatially unconstrained whole-scalp analyses. Applying the conventional cluster-level threshold of *p <* 0.05 did not change any inferential result.

### Population-state geometry and dynamics

#### Inversion and common-mode decomposition

The Learning-minus-Sham aperiodic slope effects for each participant were averaged across the frontal ROI and denoted Δ*_W,i_* for pre-sleep Wake and Δ*_N,i_* for early NREM. Paired effects were transformed into orthogonal inversion and common-mode coordinates:

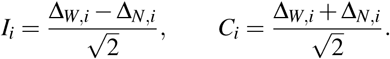

The inversion coordinate quantified opposite-direction changes across Wake and NREM, whereas the common-mode coordinate quantified changes in the same direction across both states. Positive inversion values corresponded to the learning-related pattern of flatter slopes during Wake and steeper slopes during NREM. At each electrode, the corresponding cross-state contrast was defined as Δ*_W,i,e_* Δ*_N,i,e_* and tested against zero using the sensor-space cluster-based permutation procedure described above.

The group-level pattern was summarized by the centroid (*Ī*, *C̄*) and its angular deviation from the positive inversion axis, *φ* = atan2(*C̄*, *Ī*), such that *φ* = 0° indicated perfect alignment with the flatter-Wake, steeper-NREM inversion direction. Participant-level inversion dominance was quantified as |*I_i_| − |C_i_*|, with positive values indicating stronger inversion-than common-mode expression. Uncertainty in the group centroid and angular alignment was estimated by participant bootstrap resampling as described below. The mean inversion coordinate was tested against zero using a two-sided participant sign-flip test, whereas inversion dominance was tested using a one-sided sign-flip test.

#### Distributional analyses of aperiodic slope

Distributional analyses used the window-resolved 1/f slope estimates from the same participants and state-specific intervals as the primary spectral analyses. Within each 3-min window, slope estimates were averaged across the frontal ROI, and Learning and Sham distributions were constructed separately for each participant. Participant-level empirical cumulative distribution functions (ECDFs) were averaged across participants so that each participant contributed equally regardless of the number of available windows. Learning and Sham distributions were compared within participants using a signed Kolmogorov–Smirnov statistic, defined as the ECDF difference at the point of maximum absolute separation. Negative values indicated a shift toward flatter slopes and positive values a shift toward steeper slopes. Group effects were tested using participant sign-flip permutation tests as described below.

For supplementary electrode-level analyses in Fig. 9.1, distributions were constructed separately at each electrode and summarized using first-order Wasserstein distance and the signed area under the Learning-minus-Sham ECDF-difference curve (sAUC). Electrode-wise sAUC tests were corrected for multiple comparisons using the FDR procedure.

#### Cross-fitted population geometry

Population geometry was constructed from window-resolved aperiodic slope estimates across the 121 scalp electrodes. Within each Participant State Electrode combination, Learning and Sham windows were pooled and robustly standardized using the pooled median and median absolute deviation. Participant-level learning-effect vectors were then calculated as the difference between the mean standardized Learning and Sham population vectors. Analyses requiring both Wake and NREM learning-effect maps included 23 participants.

State-specific population axes were estimated using leave-one-participant-out (LOPO) cross-fitting. For each held-out participant, the population axis was defined as the unit-normalized mean learning-effect vector of the remaining participants, and the held-out participant’s effect was projected onto this independently estimated axis. Wake and NREM axes were estimated separately using all participants available within each state. For participants with both Wake and NREM data, a reciprocal population axis was defined for each held-out participant as the unit-normalized difference between the mean Wake and NREM learning-effect vectors of the remaining participants. The axis was oriented so that the training-sample Wake effect had a positive projection. The held-out participant’s Wake and NREM projections were denoted *u_i_* and *v_i_*, respectively, and reciprocal strength was defined as 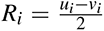, such that positive values indicated stronger expression of the Wake-positive/NREM-opposing population pattern. State-specific and reciprocal-axis inference used participant-level condition-label sign-flip permutations, with the relevant LOPO axis rebuilt on every permutation; participant-bootstrap confidence intervals reconstructed the same cross-fitting procedure. Spatial-pattern reproducibility was evaluated in supplementary analyses using within-participant split-half validation and LOPO cross-participant generalization (Fig. S10.1).

Learning and Sham windows from each held-out participant were also projected onto the reciprocal axis to characterize distribution-wide displacement. At each of the 10th–90th percentiles, displacement was defined as the Learning-minus-Sham difference in the corresponding projected quantile. The mean displacement across the nine quantiles provided a single participant-level summary for inference.

Changes in population-state occupancy were quantified by dividing the pooled Learning and Sham projections within each Participant × State combination into five equal-probability bins and calculating the Learning-minus-Sham occupancy difference in each bin. Because the five occupancy differences sum to zero, whole-profile redistribution was quantified by the Euclidean norm of the group-mean occupancy vector,

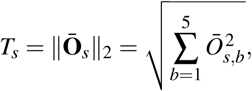

and tested against the upper tail of the full-refit condition-label permutation distribution. For two-dimensional visualization, the reciprocal axis was used as the first dimension, and the leading principal component of the residual orthogonal activity as the second dimension. Condition-specific occupancy densities were normalized within participant before averaging across participants. The whole-profile occupancy statistic was tested against the upper tail of a full-refit condition-label permutation distribution.

#### Spatial organization of reciprocal population geometry

To distinguish regional structure from the global Wake–NREM shift, each participant’s Wake and NREM Learning-minus-Sham maps were centered by their respective whole-scalp means. The residual reciprocal map was then defined as the centered Wake effect minus the centered NREM effect, 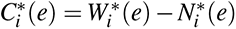 such that positive values indicated stronger reciprocal expression than that participant’s whole-scalp average.

Cross-state spatial correspondence was quantified by correlating the group-mean global-adjusted Wake map, *W^∗^*(*e*), with the group-mean sign-reversed NREM map, *N^∗^*(*e*), across the 121 electrodes using Spearman rank correlation. Regional organization was assessed by centering the five ROI means by their electrode-count-weighted mean and testing the sum of squared standardized ROI contrasts using joint participant sign flips applied to the complete five-dimensional profile. The a priori frontocentral contrast was quantified as the mean residual reciprocal effect across the combined frontal and central ROIs (50 electrodes) minus the mean across the remaining 71 electrodes. An anatomical null compared this contrast with equally sized randomly reassigned electrode sets. Wake and opposing-NREM contributions to the frontocentral effect were also tested separately. Evidence for contributions from both states was summarized by the conjunction statistic *P*_conjunction_ = max(*P_W_, P_−N_*). The frontocentral enrichment contrast, its anatomical null, and the separate Wake and opposing-NREM enrichment tests were tested one-sided.

#### Temporal evolution of reciprocal population geometry

Learning-minus-Sham multichannel effects were calculated at relative times available in both conditions for each participant and projected onto the participant-specific reciprocal LOPO axis to obtain the time-resolved expression of the reciprocal population pattern. To assess cross-state temporal correspondence, early and late Learning-minus-Sham maps were calculated from the first and final 20% of eligible Wake and NREM windows. NREM maps were sign-reversed so that positive values indicated expression opposing the Wake perturbation. Correspondence was quantified for the four Wake–NREM pairings (early Wake–early NREM, early Wake–late NREM, late Wake–early NREM and late Wake–late NREM). For each ordered pair of phase maps *A* and *B*, the LOPO source axis for participant *i* was the unit-normalized mean *A* map from all other participants,

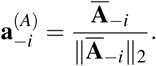

Directional correspondence from *A* to *B* was the signed projection

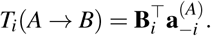

Pairwise cross-state correspondence was defined as

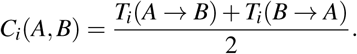

Temporal change within each state was quantified separately in participants contributing at least 10 min of data in both conditions, comprising 11 participants for Wake and 25 for NREM. Learning-minus-Sham projections were calculated from the median projection in the first and final 20% of eligible windows, and the temporal effect was defined as the late-minus-early difference. Early-to-late effects were tested separately in Wake and NREM using two-sided participant sign-flip tests, with the reciprocal axes rebuilt on every permutation. The analysis was repeated using 10%, 20%, 33% and 50% temporal-edge definitions as a sensitivity analysis.

#### Wake-defined restorative NREM trajectories

To test whether NREM dynamics were specifically organized with respect to the preceding Wake perturbation, a Wake-only population axis was constructed independently for each NREM participant from the mean Learning-minus-Sham Wake effect of the remaining eligible Wake participants. The Wake-defined reference axis was estimated from participants who contributed at least 10 min of usable pre-sleep Wake data in both Learning and Sham to provide a temporally stable estimate of the waking learning perturbation. As a sensitivity analysis, the complete procedure was repeated using all 26 participants with paired Wake recordings, which preserved the significant Learning-related restoration effect in all trajectory analyses described below. No NREM learning effect was used to define or orient this axis. NREM Learning and Sham recordings were relabelled as anonymous nights, and condition identity remained concealed during trajectory detection. The two anonymous NREM recordings for each participant were truncated to their common available duration and projected onto the Wake-defined axis. Trajectory points were sampled at 3-min intervals, and all contiguous intervals spanning at least three sampled points were evaluated. For each candidate interval containing projected values *q*_1_*, . . ., q_m_* at times *t*_1_*, . . ., t_m_*, a linear temporal slope *β*^^^ was estimated. Because positive projection values indicated expression in the Wake-perturbation direction, endpoint displacement was defined as *D* = *q*_1_ *q_m_*, such that *D >* 0 indicated net motion opposite to the Wake perturbation. Restoration strength was defined as

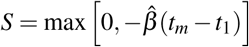

for intervals with *β*^^^ *<* 0 and *D >* 0, and as zero otherwise. Each candidate interval contributed one restoration-strength value. After all intervals had been scored while condition labels remained concealed, the anonymous recordings were mapped back to Learning and Sham. All interval-level restoration-strength values were retained, and the Learning and Sham distributions were compared within participant using a tie-corrected probability-of-superiority AUC. Candidate intervals could overlap, but inference was reduced to one AUC per participant, with 0.5 indicating no condition difference.

Direction specificity was tested using fixed-duration trajectories spanning 9, 12, 15 and 18 min. Motion opposite to and toward the Wake perturbation was separated into restorative and anti-restorative components, and Learning-versus-Sham AUCs were calculated for each direction. Direction specificity was defined as the restorative minus anti-restorative AUC and summarized across trajectory durations.

An exact Wake-source label test assessed whether restoration depended specifically on the experimentally defined Wake learning geometry. All possible Learning/Sham label assignments among the Wake source participants were enumerated, with the Wake-only axis and subsequent restoration analysis reconstructed for each assignment. The observed mean participant AUC was tested against the upper tail of this exact randomization distribution. To control for regression to the mean, Learning and Sham NREM trajectories were matched within participant for both their initial position along the Wake-defined axis and their timing within NREM. Wake-opposing motion was then compared between conditions using the matched trajectories.

For the perturbation-restoration analysis, each participant’s Wake perturbation was defined as the signed projection of their Wake Learning-minus-Sham effect onto a Wake-only LOPO axis estimated from the remaining participants. Static NREM restoration was the sign-reversed projection of that participant’s NREM effect onto the same axis. Wake perturbation was related separately to static NREM restoration and to the participant-level Learning-versus-Sham restoration AUC.

### Neural–behavioral analyses

Neural–behavioral analyses included the 22 participants with complete behavioral, pre-sleep Wake and NREM data. The primary behavioral outcome was the sleep-specific memory benefit defined above. Participant-level neural predictors were derived from Learning-minus-Sham effects. Pre-sleep Wake and NREM aperiodic slope composites were calculated from the electrodes belonging to the corresponding significant sensor-space clusters identified in the neural analyses. The same inversion and common-mode decomposition described in the geometry analysis was applied to these cluster-derived Wake and NREM composites. Pre-sleep theta, NREM SWA and NREM sigma composites were calculated across all 121 scalp electrodes since no significant cluster was identified. For electrode-based composites, values at each electrode were standardized across participants before averaging across electrodes within participant. Neural feature construction was independent of behavioral outcome.

The cross-state geometry score quantified participant-level expression of the reciprocal Wake-to-NREM population pattern defined above. For each participant, the reciprocal axis was estimated from the neural data of the remaining participants, and the held-out participant’s Wake and NREM learning-effect maps were projected onto this independently estimated axis. Their signed Wake-minus-NREM projection contrast defined the geometry score. Associations of sleep-specific memory benefit with the Wake slope, NREM slope, static inversion and geometry score were assessed using Spearman correlations. Partial-correlation inference used rank residualization with Freedman–Lane permutation. Robustness of the neural–memory association was assessed by repeating the analysis after omitting each participant in turn.

To characterize the multivariate neural relationship with memory, we applied partial least squares correlation (PLSC)^83^ to six neural predictors: cross-state geometry score, static inversion, common mode, pre-sleep theta, NREM SWA and NREM sigma. Cross-state geometry, theta, SWA and sigma were standardized across participants before PLSC. Static inversion and common mode were constructed from standardized Wake and NREM slope composites using the orthonormal transformation described above and were not re-standardized thereafter. The behavioral vector contained standardized sleep-specific memory benefit. Wake and NREM aperiodic slope measures were not included as fitted predictors because their information was already represented by the inversion and common-mode coordinates. Instead, they were projected onto the fitted latent axis as supplementary state-specific measures. The first neural and behavioral latent scores were obtained from the corresponding PLSC projections. The sign of the first latent mode was oriented so that the behavioral latent score was positively aligned with memory benefit. Significance of the first latent mode was assessed by permutation of the behavioral labels and recomputation of the singular value.

Neural structure correlations were calculated as Pearson correlations between each neural measure and the first neural latent score, including the supplementary Wake and NREM projections. For bootstrap inference, PLSC was refit within each resample and sign-aligned to the full-sample solution before structure correlations were recomputed. Stability of the neural axis was assessed by LOPO refitting; each refitted axis was sign-aligned to the full-sample solution before cosine similarity was calculated. Feature-family preservation was examined by separately refitting PLSC with the three aperiodic measures (geometry, static inversion and common mode) and the three oscillatory measures (theta, SWA and sigma). Each reduced neural axis was embedded in the original six-dimensional feature space by assigning zero weights to omitted features, sign-aligned to the full solution, and compared with the full neural axis using cosine similarity.

Held-out prediction of sleep-specific memory benefit was tested using outer LOPO cross-validation. Five single-predictor models were tested on the same folds: pre-sleep Wake slope, NREM slope, static inversion, cross-state geometry score and common mode. An oscillatory model containing pre-sleep theta, NREM SWA and NREM sigma was tested using the same outer LOPO procedure as a separate benchmark. Predictors were standardized using the training participants only, and static inversion and common mode were reconstructed within each training fold. Because the geometry score depends on a population-level reciprocal axis, its predictive analysis was nested within the outer LOPO procedure. Within each outer fold, the geometry score for every training participant was itself cross-fitted using a reciprocal axis estimated from the neural source pool after excluding both that participant and the outer test participant. The outer test participant was scored using an axis estimated without that participant. Thus, the outer test participant contributed neither to the behavioral regression nor to any reciprocal axis used to construct the training predictors.

Held-out predictive performance was quantified as

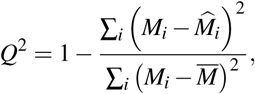

where *M_i_* is the observed sleep-specific memory benefit, *M^i* is its LOPO prediction and *M* is the sample mean. Positive *Q* permuting behavioral outcomes and repeating the LOPO procedure. Participant-bootstrap distributions of *Q*^2^ were obtained by resampling paired observed outcomes and held-out predictions, with 95% BCa confidence intervals calculated from the corresponding jackknife estimates. Permutation *P* values for the five single-predictor models were corrected together using FDR.

Prespecified differences in held-out *Q*^2^, including the comparison between cross-state geometry and other simpler models, were tested by sign-flipping participant-level differences in squared prediction error, with multiplicity across these comparisons controlled using FDR. As a behavioral-specificity analysis, neural predictors were each related to sleep-specific retrieval-latency change, defined as the Sleep-minus-Wake difference in delayed-minus-immediate log mean RT on correct trials (Table S13.1).

### Statistical analysis

All behavioral and neural–behavioral statistical analyses were conducted in R^84^, whereas neural data processing and neural-only statistical analyses were conducted in Python. Unless otherwise specified, statistical tests were two-sided, permutation tests used 10,000 iterations, and confidence intervals were estimated from 10,000 participant-level bootstrap resamples. Permutation and test statistics were defined according to the structure of each analysis as described above. Participant-level one-sample and paired contrasts were tested using sign-flip permutation tests, whereas the cross-state topographic correspondence used within-participant electrode-label permutations. Residual-permutation analyses used the Freedman–Lane procedure^85^. Correlation analyses used Spearman rank coefficients, except for PLSC latent score and structure correlations, which used Pearson coefficients. Bootstrap confidence intervals were calculated using bias-corrected and accelerated (BCa) intervals with leave-one-participant-out jackknife acceleration. Generalized linear mixed-model estimated marginal means and model-derived contrasts used Wald confidence intervals computed on the model scale and transformed to the response scale where appropriate. Continuous predictors in mixed-effects models were mean-centered before fitting, and nested models were compared using likelihood-ratio tests where specified. Outliers were identified using the median absolute deviation (MAD) criterion with a threshold of 3.5^86^. Primary statistics were calculated from the full analysis samples; analyses excluding identified outliers were used as sensitivity checks. Multiple comparisons were controlled within prespecified families using the Benjamini–Hochberg false discovery rate procedure (*q* = 0.05) unless otherwise specified. Sensor-space cluster analyses used cluster-level family-wise error control, and the three prespecified correlation tests in Fig. 4d,g,j were Bonferroni-corrected. All bootstrap confidence intervals, effect sizes, test statistics, and exact *P* values are reported in the public repository as described below.

## Data availability

All behavioral data and preprocessed neural data used in the reported analyses are available for replication through the Open Science Framework repository: https://osf.io/y43we/. The raw high-density EEG dataset exceeds 3.4 TB and is maintained on local institutional storage. In addition, participants were not asked to provide consent for public data sharing. The raw data are available from the corresponding author upon reasonable request.

## Code availability

Code for data analysis and visualization is available in the Open Science Framework repository, https://osf.io/y43we/.

## Author contributions

Conceptualization: T.N., M.B. and R.M.C.S. Experimental design: M.B. and R.M.C.S. Data collection: M.B., with contributions from L.C., H.P. and N.V. Data preprocessing: T.N., with contributions from M.B. Analytical methodology and code:

T.N. Formal analysis: T.N., with initial analyses by M.B. and input from A.A. Visualization: T.N. Manuscript preparation: T.N., with contributions from M.B. Manuscript revision: T.N. and R.M.C.S. Project supervision and administration: R.M.C.S. Funding acquisition: R.M.C.S.

