## Supplementary material for "Sleep renormalizes learning-perturbed cortical dynamics to stabilize memory"

Thea Ng[1,2]*, Morgan Barnes[3]*, Atif Abedeen[4], Lucas Collignon[5], Hirni Patel[6,7], Nicholas Vovcsko[7], Rebecca M. C. Spencer[3,7,8]†

#### Contents

**1 Participant characteristics and data inclusion** **2**

**2 Sleep architecture and NREM depth** **4**

**3 Regions of interest electrode assignment** **5**

**4 *Specparam* fit-quality diagnostics** **7**

**5 Electrode-level spectral estimate quality control** **8**

**6 Robustness of 1/f slope effects across frequency range and reference** **15**

**7 Robustness of 1/f slope effects across spectral fitting approaches** **17**

**8 Robustness of 1/f slope effects to the analysis window** **18**

**9 Electrode-level epoch-wise distribution of 1/f slopes** **19**

**10 Cross-participant and cross-epoch generalization of aperiodic patterns** **20**

**11 REM sleep aperiodic slope analysis** **21**

**12 REM aperiodic slope and memory association** **22**

**13 Neural associations with retrieval-latency change** **23**

*[1]Neuroscience & Behavior Program, Mount Holyoke College. [2]Department of Mathematics & Statistics, Mount Holyoke College. [3]Neuroscience & Behavior Program, University of Massachusetts Amherst. [4]Manning College of Information and Computer Science, University of Massachusetts Amherst. [5]Department of Biology, University of Massachusetts Amherst. [6]Department of Biochemistry and Molecular Biology, University of Massachusetts Amherst. [7]Department of Psychological & Brain Sciences, University of Massachusetts Amherst. [8]Institute of Applied Life Sciences, University of Massachusetts Amherst. *These authors contributed equally to this work. † Correspondence:

### 1 Participant characteristics and data inclusion

| | *N* | Mean (s.d.) |
|---|---|---|
| Total sample | 36 | |
| *Racial identity* | | |
| White | 20 | |
| Asian | 13 | |
| Black | 1 | |
| More than one racial identity | 1 | |
| Missing | 1 | |
| *Gender* | | |
| Female | 20 | |
| Male | 15 | |
| Missing | 1 | |
| Age, years | | 23.37 (1.99) |
| Insomnia Severity Index | 35 | 4.31 (3.45) |
| Morningness-Eveningness Questionnaire | 35 | 47.80 (8.95) |
| Pittsburgh Sleep Quality Index | 35 | 5.20 (2.59) |
| *Stanford Sleepiness Scale* | | |
| Learning overnight session, encoding | 32 | 2.84 (1.22) |
| Learning overnight session, recall | 32 | 2.12 (1.16) |
| Wake session, encoding | 27 | 2.22 (0.80) |
| Wake session, recall | 22 | 1.82 (0.80) |
| Sham session, encoding | 30 | 2.97 (1.16) |
| Sham session, recall | 29 | 1.97 (0.87) |

**Table S1.1 — Sample characteristics and questionnaire data.**

Notes. Means and standard deviations of demographic variables and questionnaire measures are reported for all collected samples. Values are reported as counts or mean (s.d.). s.d., standard deviation; ISI, Insomnia Severity Index; MEQ, Morningness-Eveningness Questionnaire; PSQI, Pittsburgh Sleep Quality Index; SSS, Stanford Sleepiness Scale. PSQI scores range from 0 to 21, with scores $>6$ indicating poor sleep quality. MEQ scores range from 16 to 86, with scores $>59$ indicating morning type and scores $\leq 41$ indicating evening type. ISI scores range from 0 to 28, with scores $\geq 15$ indicating clinical insomnia. SSS scores range from 1 to 8, where 1 indicates fully awake and 8 indicates asleep. ISI scores indicated that participants were free from insomnia symptoms. On average, MEQ scores indicated an intermediate circadian preference, and no participants were extreme morning or evening types. PSQI responses indicated typical sleep quality for young adults, with a sample mean below 6.

| Sample or analysis | Final $N$ | Appears in | Inclusion basis or primary reason for reduced sample size |
|---|---|---|---|
| Enrolled participants | 41 | Methods | All participants enrolled in the study. |
| Included study cohort | 36 | Methods; Table S1.1 | Five enrolled participants were excluded due to missing behavioral data and unusable EEG data. |
| Behavioral analysis sample | 34 | Fig. 2 | Two participants had missing post-sleep recall data. |
| *Single-metric PSG analyses* | | | |
| – Pre-sleep wake 1/f slope | 26 | Figs. 3, 4, 5 | Unusable or irregular PSG data, unsuccessful spectral parameterization, or insufficient usable pre-sleep wake EEG duration in one or both sessions |
| – Pre-sleep wake 1/f temporal analysis | 11 | Fig. 5 | At least 10 min of usable data was required in both Learning and Sham to provide sufficient temporal coverage and stability for within-state temporal comparisons. |
| – Pre-sleep wake theta power | 26 | Fig. 4 | Restricted to the same valid paired pre-sleep wake sample used for the 1/f slope analysis. |
| – NREM 1/f slope | 25 | Figs. 3, 4, 5 | Unusable or irregular PSG data, or unsuccessful spectral parameterization. |
| – NREM slow-wave activity (SWA) | 25 | Fig. 4 | Restricted to the same valid paired NREM sample used for the 1/f slope analysis. |
| – NREM sigma power | 25 | Fig. 4 | Restricted to the same valid paired NREM sample used for the 1/f slope analysis. |
| – REM 1/f slope | 26 | Fig. S11.1 | Unusable or irregular PSG data, or unsuccessful spectral parameterization. |
| – Post-sleep wake 1/f slope | 6 | Fig. 3 | Post-sleep wake EEG was not consistently acquired following morning awakening because recordings were not systematically continued before cap removal. At least 5 min of usable post-sleep wake EEG was required in both the learning and sham sessions, unusable or irregular PSG data, or unsuccessful spectral parameterization. |
| *Cross-state analyses* | | | |
| – Combined Wake–NREM analyses | 23 | Figs. 3, 5 | Restricted to participants contributing valid paired Learning–Sham 1/f slope data in both pre-sleep Wake and NREM. |
| *Neural–memory analyses* | | | |
| – Multivariate neural–memory analysis | 22 | Fig. 6; Fig. S12.1; Table S13.1 | Complete-data intersection across all neural predictors and behavioral measures included in the multivariate model. |

**Table S1.2 — Analysis-specific sample sizes and reasons for reduced inclusion.**

Notes. Values are the final participant counts used in each analysis. Single-metric PSG analyses used the largest valid paired learning–sham sample available for the corresponding state and measure and were not restricted to a common participant intersection.

### 2 Sleep architecture and NREM depth

| | Learning Mean (s.d.) % of TST | Sham Mean (s.d.) % of TST | *p*-value |
|---|---|---|---|
| TST | 396.3 (78.93) | 405.5 (76.71) | 0.62 |
| SOL | 33.9 (35.31)<br>8.55% | 29.9 (25.94)<br>7.37% | 0.59 |
| WASO | 22.41 (27.73)<br>5.65% | 15.45 (21.09)<br>3.81% | 0.24 |
| NREM 1 | 13.77 (10.92)<br>3.47% | 12.53 (7.95)<br>3.09% | 0.59 |
| NREM 2 | 205.17 (68.6)<br>51.77% | 211.13 (46.5)<br>52.06% | 0.67 |
| NREM 3 | 116.24 (40.26)<br>29.33% | 112.74 (33.09)<br>27.80% | 0.69 |
| REM | 56 (28.6)<br>14.13% | 64.4 (32.8)<br>15.88% | 0.26 |

**Table S2.1 — Sleep stage characteristics.**

Notes. Average sleep stage characteristics are reported for the learning and sham conditions. Values are reported as mean (s.d.) in minutes, with the percentage of total sleep time reported below each value where applicable. TST, total sleep time; SOL, sleep onset latency; WASO, wake after sleep onset; NREM, non-rapid eye movement sleep; REM, rapid eye movement sleep; SWS, slow-wave sleep. *p*-values are reported from two-sided paired-samples *t*-tests comparing the learning and sham nights.

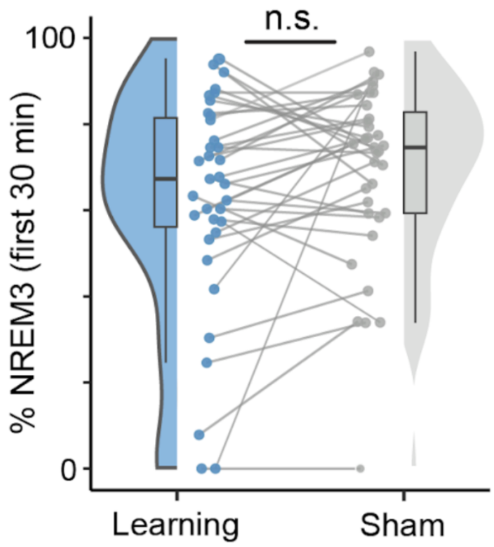


**Fig. S2.2 — NREM 3 proportion across learning and sham conditions.**

Notes. The percentage of NREM 3 sleep within the first 30 min of NREM sleep did not differ between the learning and sham conditions, indicating that the learning-related spectral effects were not attributable to differences in initial sleep depth across nights.

#### 3 Regions of interest electrode assignment

| Channel | ROI | Channel | ROI |
|---|---|---|---|
| Fp1 | Prefrontal | CCP1h | Central |
| Fp2 | Prefrontal | CCP2h | Central |
| Fpz | Prefrontal | CCP3h | Central |
| AF3 | Prefrontal | CCP4h | Central |
| AF4 | Prefrontal | CCP5h | Central |
| AFp1 | Prefrontal | CCP6h | Central |
| AFp2 | Prefrontal | P3 | Parietal |
| AF7 | Prefrontal | P4 | Parietal |
| AF8 | Prefrontal | Pz | Parietal |
| AFF1h | Prefrontal | P1 | Parietal |
| AFF2h | Prefrontal | P2 | Parietal |
| AFF5h | Prefrontal | P5 | Parietal |
| AFF6h | Prefrontal | P6 | Parietal |
| F3 | Frontal | POz | Parietal |
| F4 | Frontal | PO3 | Parietal |
| F7 | Frontal | PO4 | Parietal |
| F8 | Frontal | PO7 | Parietal |
| Fz | Frontal | PO8 | Parietal |
| F1 | Frontal | PO9 | Parietal |
| F2 | Frontal | PO10 | Parietal |
| F5 | Frontal | CPP1h | Parietal |
| F6 | Frontal | CPP2h | Parietal |
| FC1 | Frontal | CPP3h | Parietal |
| FC2 | Frontal | CPP4h | Parietal |
| FC3 | Frontal | CPP5h | Parietal |
| FC4 | Frontal | CPP6h | Parietal |
| FC5 | Frontal | PPO1h | Parietal |
| FC6 | Frontal | PPO2h | Parietal |
| FFC1h | Frontal | PPO5h | Parietal |
| FFC2h | Frontal | PPO6h | Parietal |
| FFC3h | Frontal | PPO9h | Parietal |
| FFC4h | Frontal | PPO10h | Parietal |
| FFC5h | Frontal | POO1 | Parietal |
| FFC6h | Frontal | POO2 | Parietal |
| FFT7h | Frontal | POO9h | Parietal |
| FFT8h | Frontal | POO10h | Parietal |
| FCC1h | Frontal | T7 | Temporal |
| FCC2h | Frontal | T8 | Temporal |
| FCC3h | Frontal | P7 | Temporal |
| FCC4h | Frontal | P8 | Temporal |
| FCC5h | Frontal | FT9 | Temporal |
| FCC6h | Frontal | FT10 | Temporal |
| FCz | Frontal | FT7 | Temporal |
| C3 | Central | FT8 | Temporal |
| C4 | Central | TP7 | Temporal |
| Cz | Central | TP8 | Temporal |
| C1 | Central | FTT7h | Temporal |
| C2 | Central | FTT8h | Temporal |
| C5 | Central | TTP7h | Temporal |
| C6 | Central | TTP8h | Temporal |
| CP1 | Central | TPP7h | Temporal |
| CP2 | Central | TPP8h | Temporal |
| CP3 | Central | TPP9h | Temporal |
| CP4 | Central | TPP10h | Temporal |
| CP5 | Central | P9 | Temporal |
| CP6 | Central | P10 | Temporal |
| CPz | Central | | |

**Table S3.1 — ROI electrode assignment.**

Notes. Electrodes were assigned to five anatomically motivated ROIs in a high-density EEG montage. Prefrontal electrodes included FP, AF, AFp, and AFF channels; frontal electrodes included FFC, FC, FCC, FFT, and F channels; central electrodes included C, CP, and CCP channels; parietal electrodes included P, PO, PPO, POO, and CPP channels, excluding P7, P8, P9, and P10; temporal electrodes included T, TP, FT, FTT, TTP, TPP, P7, P8, P9, and P10. O- and I-prefixed channels were not included in any ROI. M1 and M2 were excluded from the analysis channel set because they were used for linked-mastoid referencing. ROI assignments were used for participant-level regional summaries and ROI-based paired statistical tests.

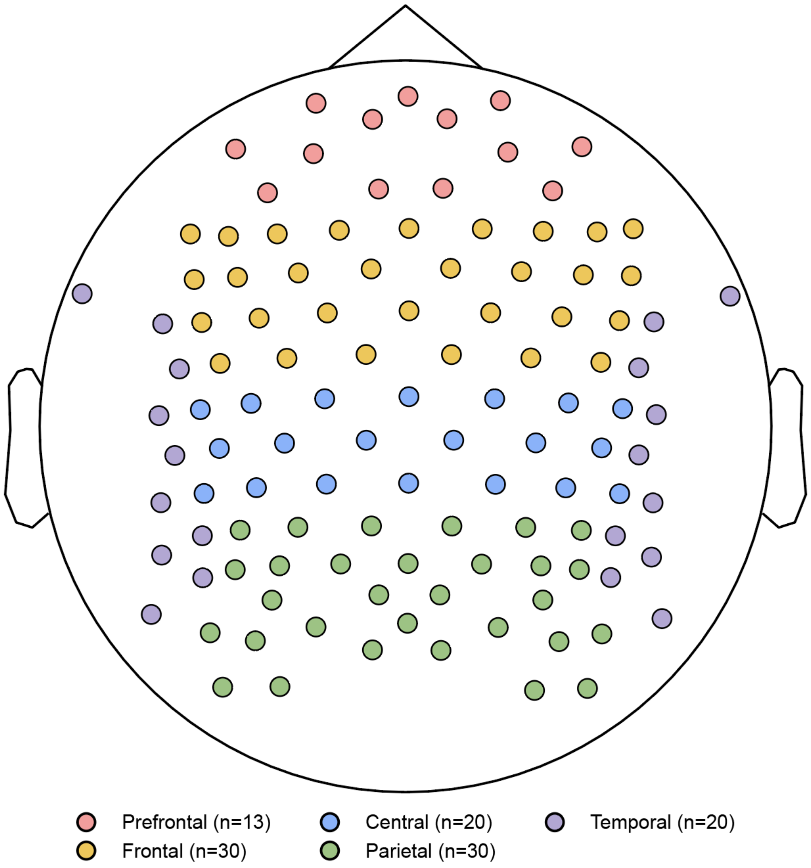


**Fig. S3.2 — ROI electrode assignment.**

Notes. Colored dots indicate the electrodes assigned to each predefined region of interest used for participant-level ROI summaries. O- and I-prefixed channels and mastoid channels were not included in ROI analyses.

### 4 *Specparam* fit-quality diagnostics

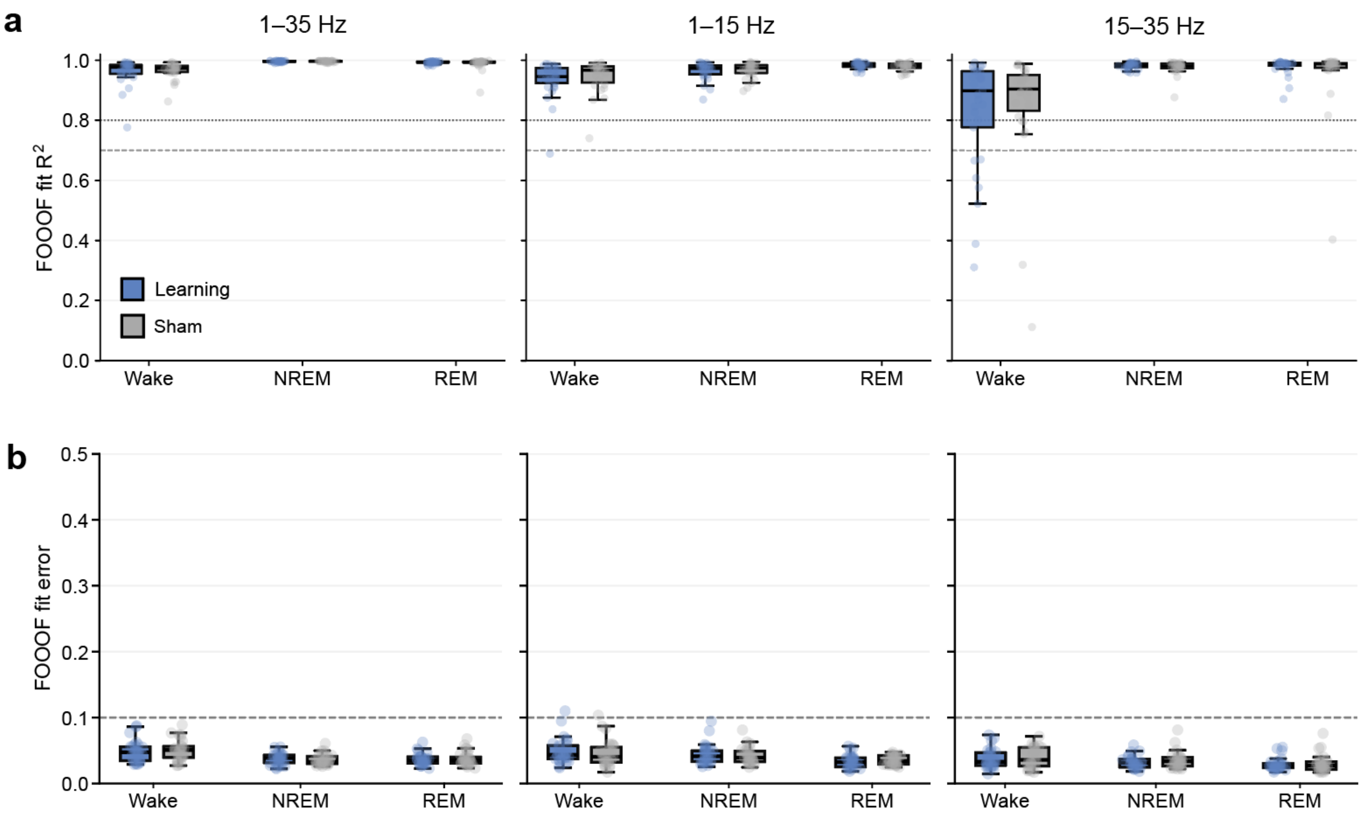


**Fig. S4.1 — *Specparam* fit-quality diagnostics across states and conditions.**

Notes. Window-level *specparam* fit diagnostics are summarized across pre-sleep wakefulness, NREM stages 2–3, and REM sleep for Learning and Sham conditions. For each participant, model $R^2$ and fit error were calculated for each eligible 3-min spectral window and then averaged across windows. Each point represents one participant-level mean of the window-resolved fit diagnostics. Columns show fits estimated over 1–35 Hz, 1–15 Hz, and 15–35 Hz. (a) Mean model $R^2$ across windows. (b) Mean model fit error across windows. Blue points and boxplots indicate Learning sessions; gray points and boxplots indicate Sham sessions. Horizontal dashed lines indicate fit-quality reference thresholds.

We next evaluated the stability of window-resolved spectral parameterization across brain state, experimental condition, and frequency fitting range. Participant-level summaries were calculated from the model $R^2$ and fit error values obtained for each eligible 3-min window. Across the 1-35 Hz fits used in the main analysis, model fit quality was consistently high across wakefulness, NREM, and REM sleep, with comparable distributions between learning and sham conditions. Similar high fit quality was observed for the 1-15 Hz fits, indicating that the lower-frequency portion of the spectrum was well captured across states and conditions. Similarly, model fit errors were uniformly low across fitting ranges and did not show systematic condition-specific differences. The 15-35 Hz fits showed greater variability in $R^2$ during pre-sleep wakefulness, but model error remained low. Together, these diagnostics indicate that the learning-related topographic effects were not attributable to broad failures of spectral parameterization or systematic differences in fit quality between learning and sham sessions.

### 5 Electrode-level spectral estimate quality control

After confirming stable spectral parameterization at the window and participant levels, we evaluated electrode-level estimates for distributional and spatial anomalies before calculating condition contrasts. Electrode-level spectral estimates were screened separately within each state, spectral parameter, and condition using robust median absolute deviation (MAD)-based modified $z$-scores,

$$z_{\mathrm{mod}} = \frac{0.6745\,(x - \mathrm{median}(x))}{\mathrm{MAD}(x)}$$

Modified $z$-scores were calculated at three complementary levels: (i) within each participant across electrodes, using $|z_{\mathrm{mod}}| \geq 4.5$; (ii) within each electrode across participants, using $|z_{\mathrm{mod}}| \geq 4.5$; and (iii) globally across all participant-by-electrode observations within the corresponding state, parameter, and condition, using $|z_{\mathrm{mod}}| \geq 6.0$. An estimate was automatically flagged if it exceeded the global threshold or simultaneously exceeded both the within-participant and within-electrode thresholds. All automatically flagged estimates were subsequently reviewed by visual inspection using the diagnostic plots shown in Figs. S5.1–S5.6. As described in the Methods, three of 32,186 visually confirmed invalid estimates were interpolated within the corresponding participant–condition–parameter electrode map using spherical splines.

All six figures use the same layout. Panels a–b show electrode-level distributions for the same representative participant across figures. Panel a shows the Learning and Sham distributions across electrodes, with dashed vertical lines indicating condition-specific medians. Panel b shows the corresponding rank-ordered electrode values. Panels c–f show group-level diagnostics. Panel c shows the group condition distributions obtained by averaging participant-specific density estimates. Panel d shows paired participant-level electrode medians. Panel e shows the distribution of Learning-minus-Sham electrode-level effects. Panel f shows the joint distribution of Learning and Sham electrode values; hexagon shading represents observation density, and black points represent participant-level medians.

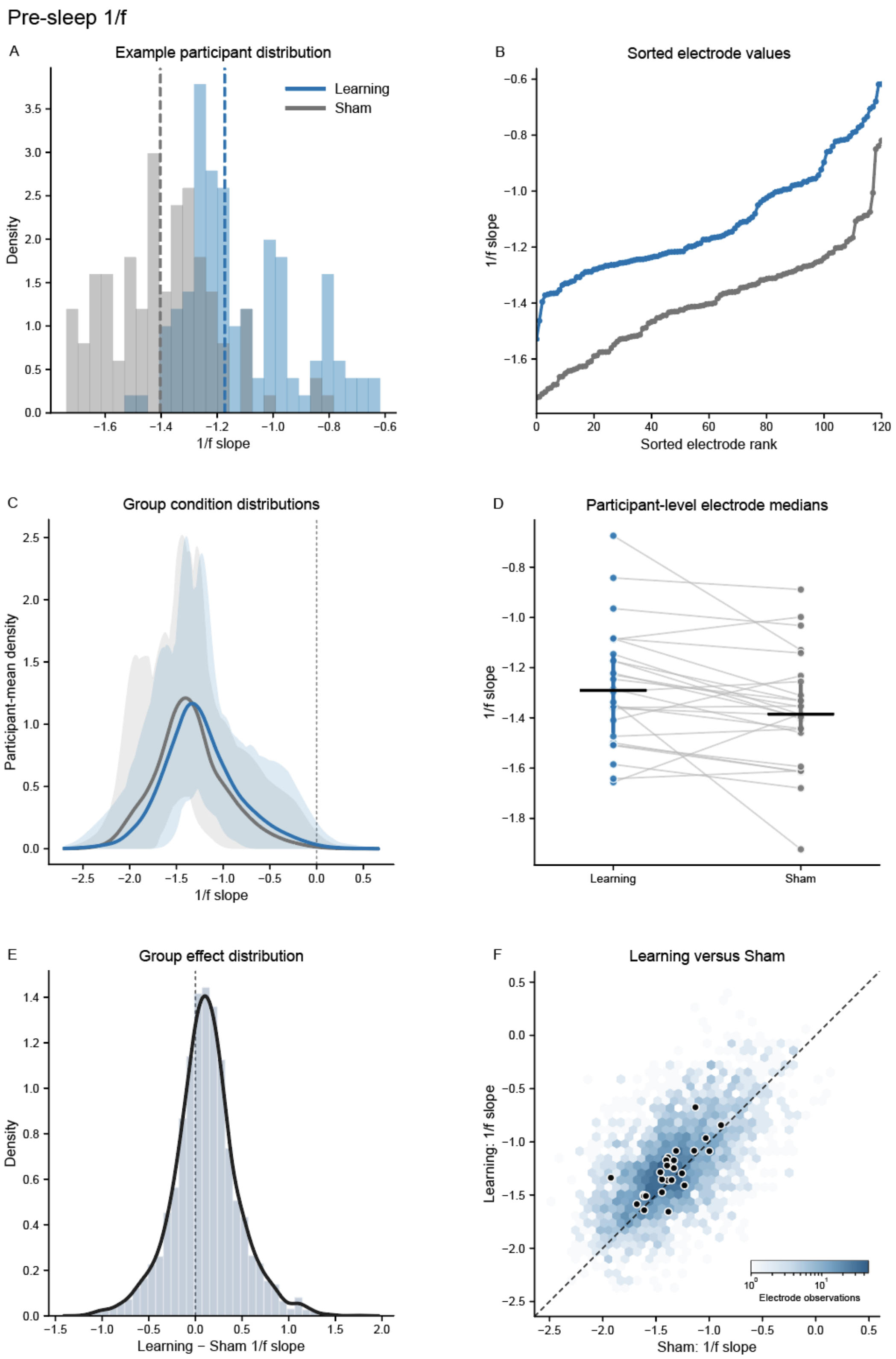


**Fig. S5.1 — Quality-control diagnostics for pre-sleep wake aperiodic 1/f slope.**

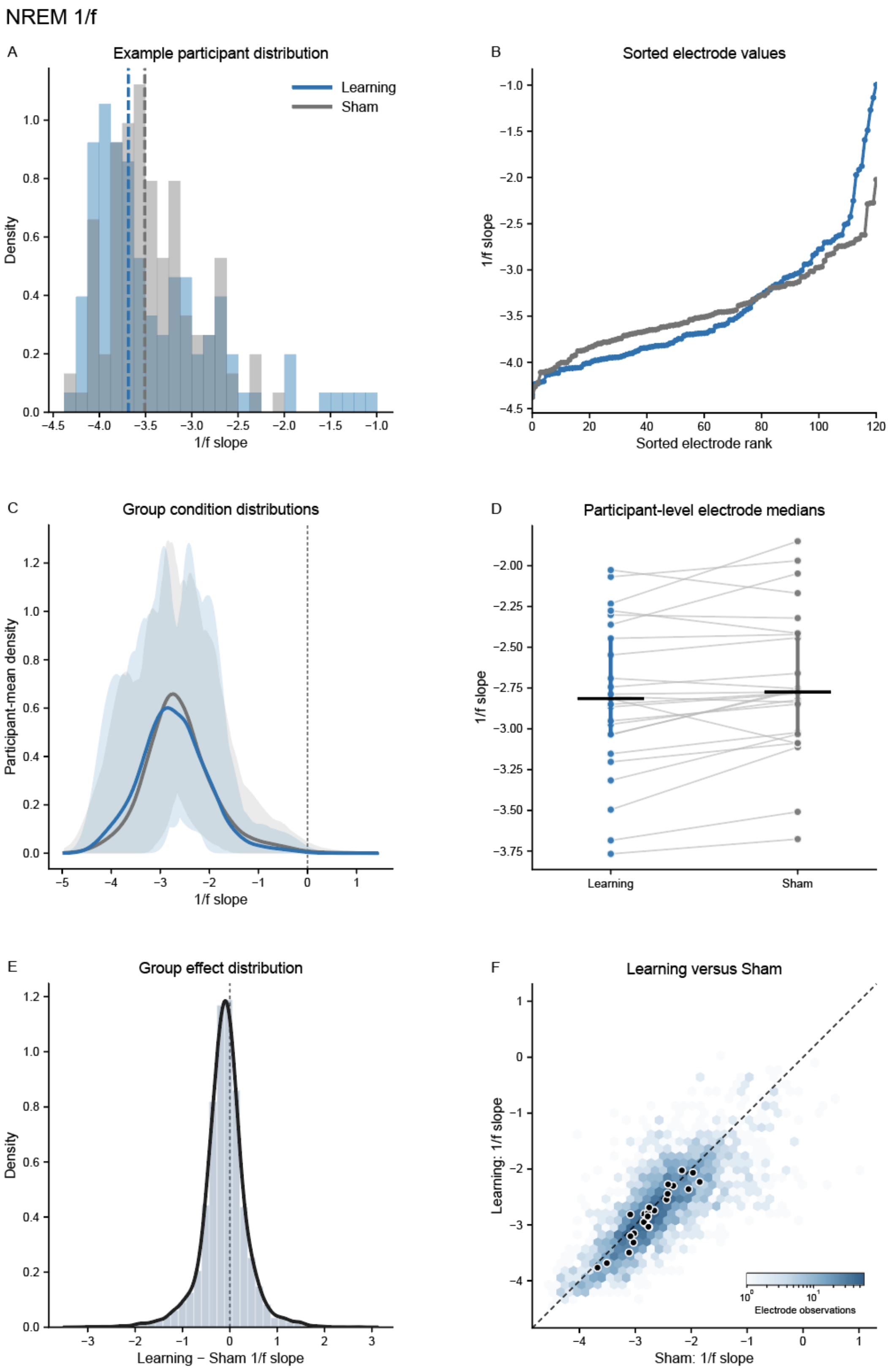


**Fig. S5.2 — Quality-control diagnostics for NREM aperiodic 1/f slope.**

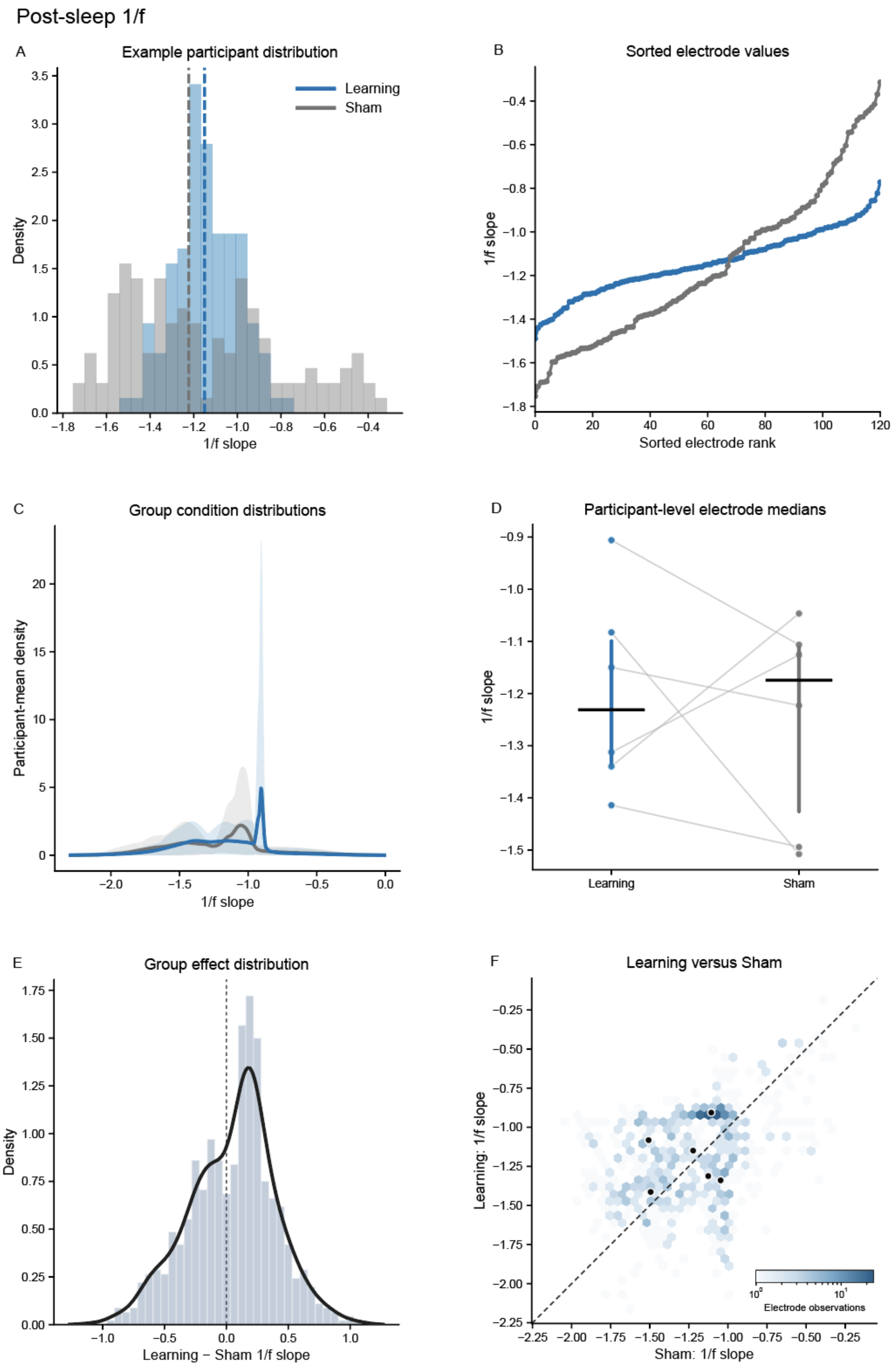


**Fig. S5.3 — Quality-control diagnostics for post-sleep wake aperiodic 1/f slope.**

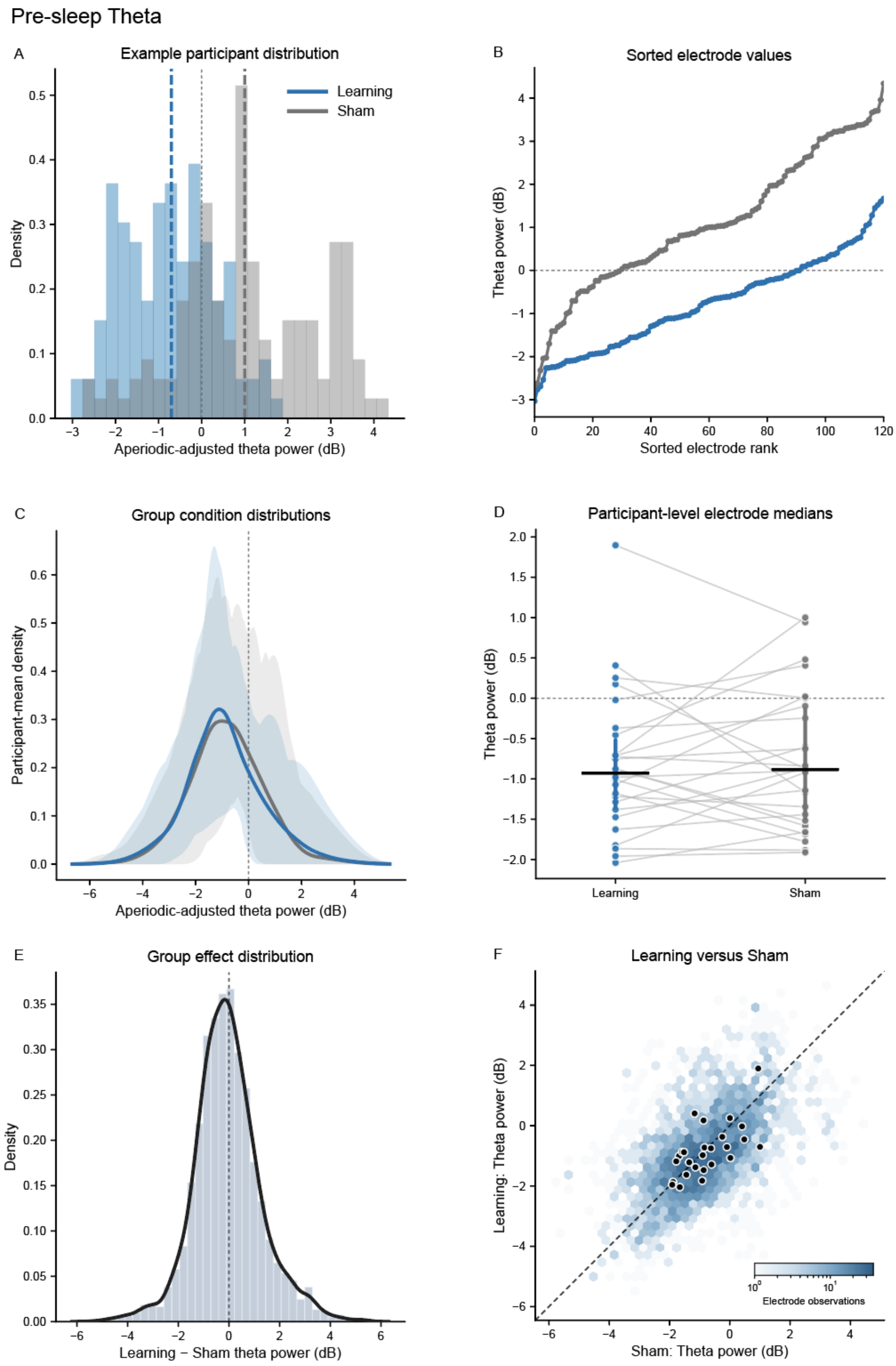


**Fig. S5.4 — Quality-control diagnostics for pre-sleep wake aperiodic-adjusted theta activity.**

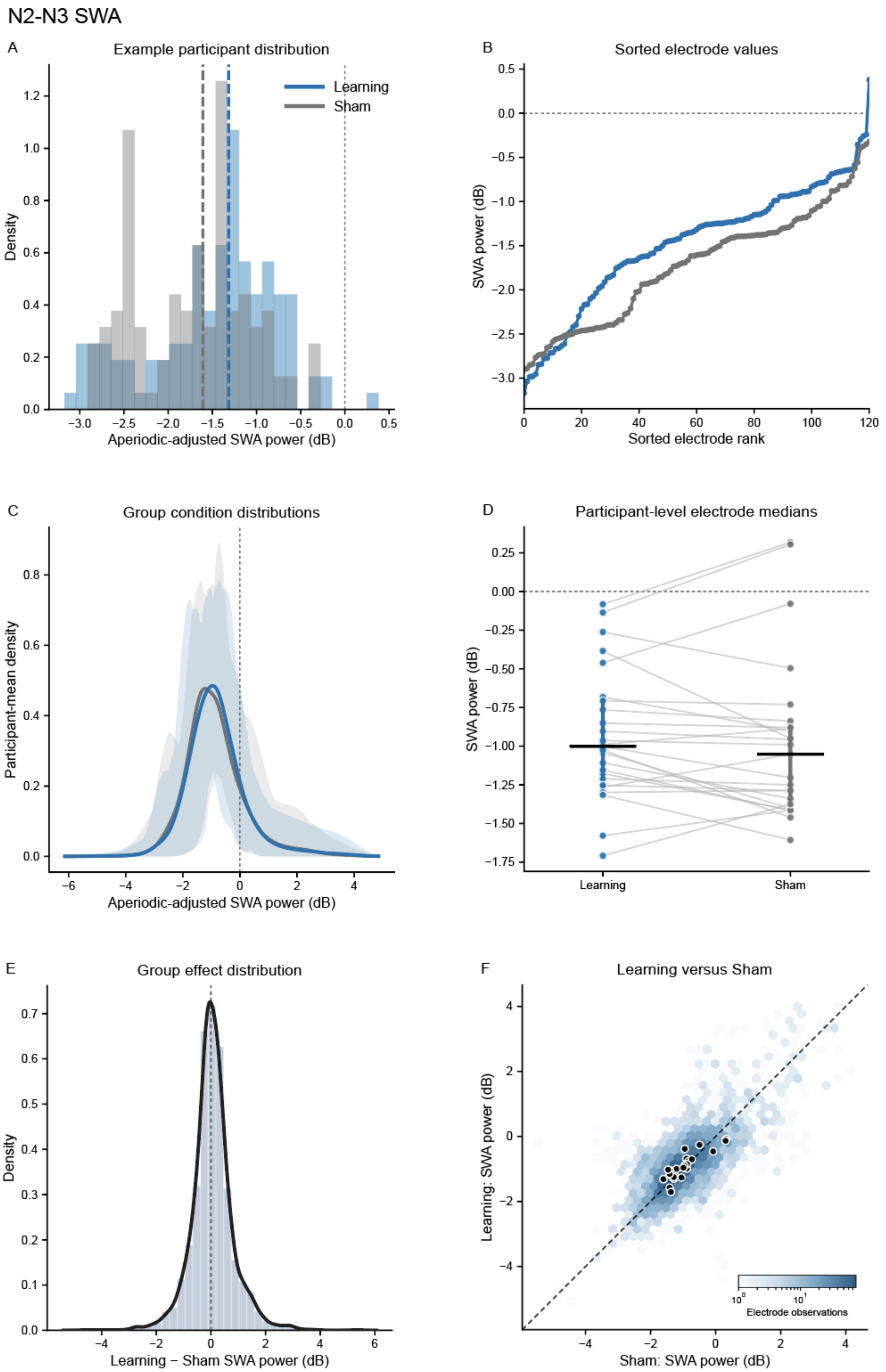


**Fig. S5.5 — Quality-control diagnostics for NREM aperiodic-adjusted slow-wave activity.**

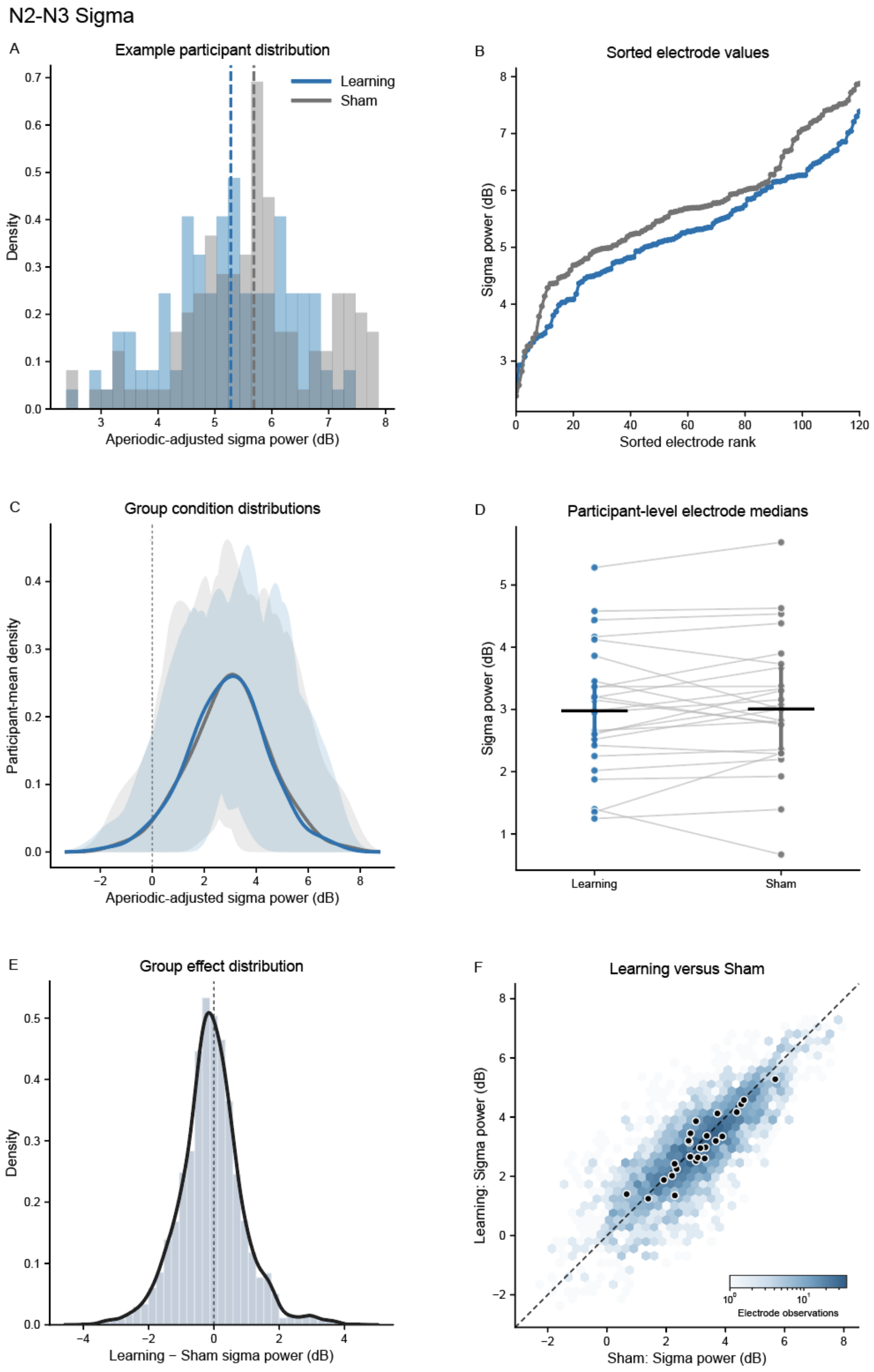


**Fig. S5.6 — Quality-control diagnostics for NREM aperiodic-adjusted sigma activity.**

### 6 Robustness of 1/f slope effects across frequency range and reference

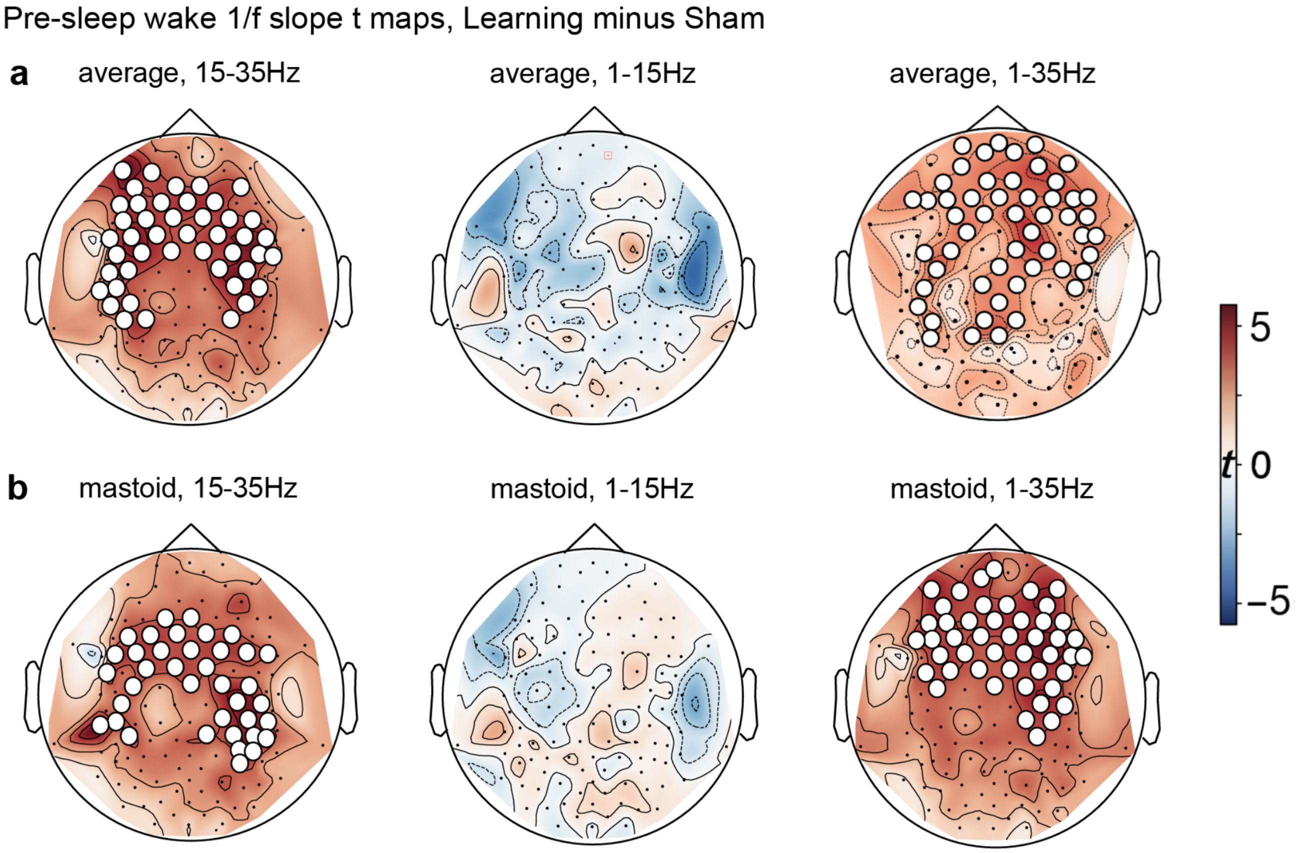


**Fig. S6.1 — Robustness of pre-sleep wake 1/f slope topographies across fitting range and reference.**

Notes. Topographic $t$-statistic maps show learning-related changes in pre-sleep wake 1/f slope, computed as Learning minus Sham, across spectral fitting frequency ranges and EEG reference channels. (a) Average-reference maps estimated over 15-35 Hz, 1-15 Hz, and 1-35 Hz after the current source density (CSD) transformation. Learning-related flattening was observed over frontal and central regions for the 15-35 Hz fit ($p_{perm} < 0.001$) and the 1-35 Hz fit ($p_{perm} = 0.002$), whereas the 1-15 Hz fit did not reproduce the frontocentral learning effect. (b) Analogous maps using mastoid reference (M1 + M2) without CSD. The positive learning-sham effect was again observed for the 15-35 Hz fit ($p_{perm} = 0.003$) and the 1-35 Hz fit ($p_{perm} < 0.001$), with no comparable frontocentral effect in the 1-15 Hz fit. White circles indicate electrodes included in significant cluster-corrected effects. Positive $t$ values indicate flatter fitted 1/f slopes in learning relative to sham condition.

To determine whether learning-related aperiodic slope topographies were robust or depended on specific frequency ranges or reference channels, we repeated the learning-sham topographic analyses across three fitting ranges and two references. During pre-sleep wakefulness, learning-related flattening was preserved across the 15-35 Hz and 1-35 Hz fits, with both average and mastoid references showing positive frontocentral effects. In contrast, the 1-15 Hz fit did not reproduce the same spatial pattern, indicating that the wake effect was primarily captured by broader or higher-frequency estimates of the aperiodic component rather than by the lower-frequency range alone.

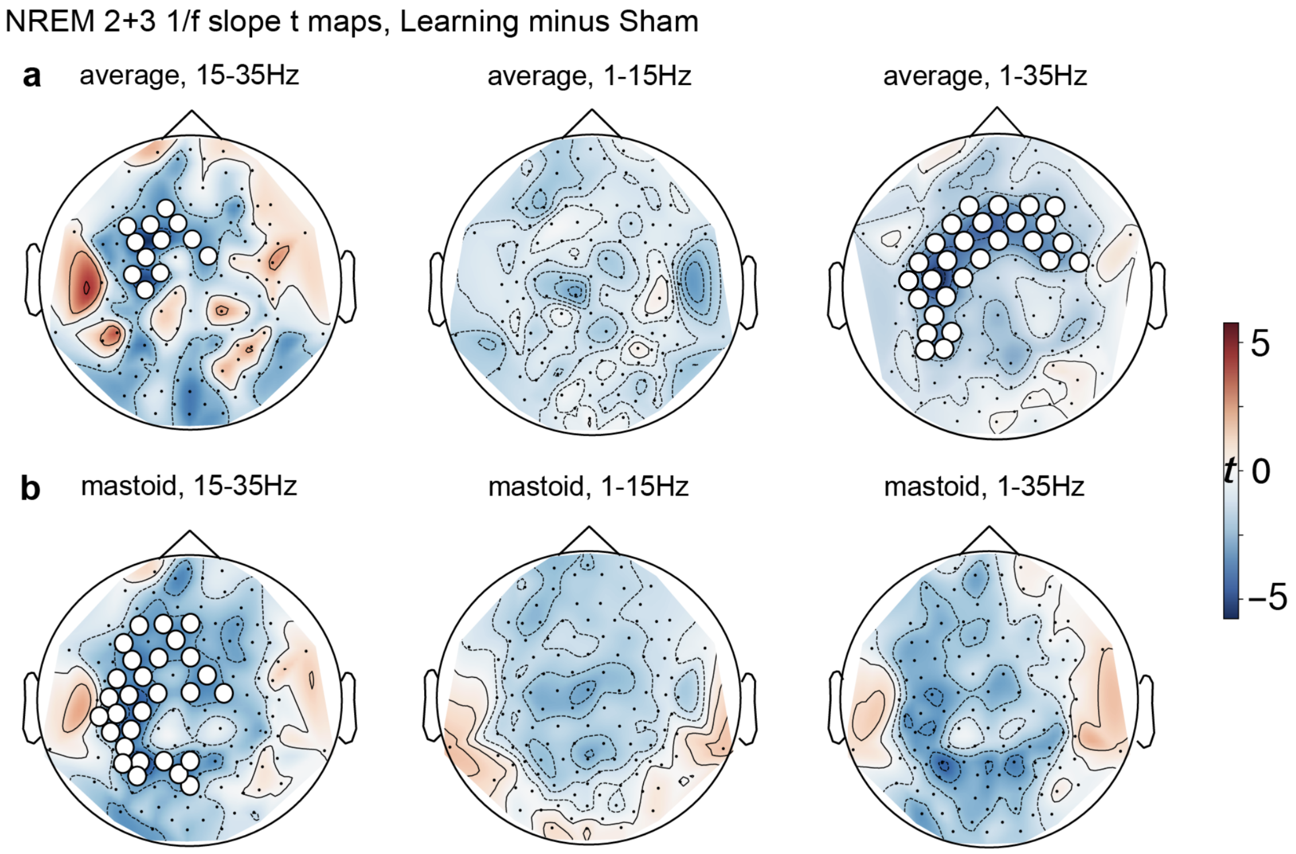


**Fig. S6.2 — Robustness of NREM 1/f slope topographies across fitting range and reference.**

Notes. Topographic $t$-statistic maps show learning-related changes in 1/f slope during the first 30 min of NREM stages 2-3, computed as Learning minus Sham, across spectral fitting frequency ranges and EEG reference channels. (a) Average-reference maps estimated over 15-35 Hz, 1-15 Hz, and 1-35 Hz after the CSD transformation. Learning-related steepening was observed over frontocentral regions for the 15-35 Hz fit ($p_{perm} = 0.037$) and the 1-35 Hz fit ($p_{perm} = 0.002$), whereas the 1-15 Hz fit did not show a comparable cluster-corrected effect (b) Analogous maps using mastoid reference (M1 + M2) without CSD. The 15-35 Hz fit showed learning-related steepening over a frontocentral distribution ($p_{perm} = 0.018$), whereas the 1-15 Hz and 1-35 Hz fits did not reach cluster-corrected significance but showed the same direction of learning-related steepening. White circles indicate electrodes included in significant cluster-corrected effects. Negative $t$ values indicate steeper fitted 1/f slopes in learning relative to sham condition.

A corresponding frequency-range dependence was observed during NREM stages 2-3. Learning-related steepening was most consistently expressed in the 15-35 Hz and 1-35 Hz fits, where the learning-induced effects emerged over frontocentral regions. The 1-15 Hz fit showed a directionally similar but non-significant pattern of learning-related steepening across both references, suggesting that the NREM effect was not driven by the lower-frequency portion of the spectrum alone. Although the exact spatial extent varied across references, the dominant pattern was preserved: learning was associated with wake flattening and subsequent NREM steepening in frontocentral scalp regions, with the effects consistent and strong across broad frequency ranges.

### 7 Robustness of 1/f slope effects across spectral fitting approaches

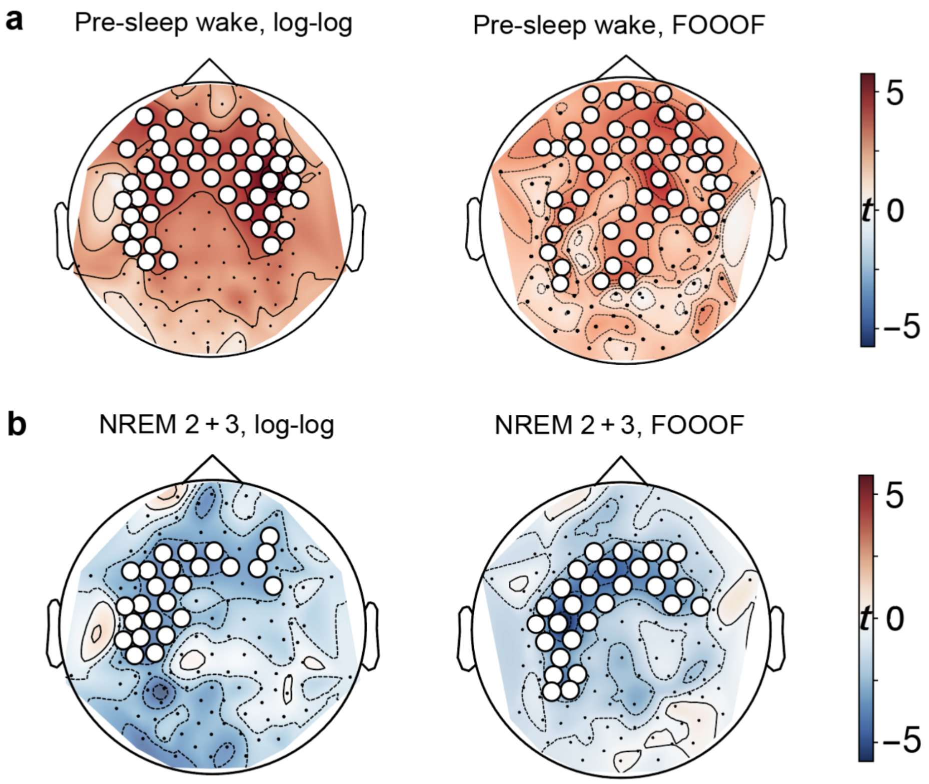


**Fig. S7.1 — Robustness of NREM 1/f slope topographies across spectral fitting approaches**

Notes. Topographic $t$-statistic maps show learning-related changes in aperiodic 1/f slope, computed as Learning minus Sham, during pre-sleep wakefulness and the first 30 min of NREM stages 2-3. (a) Pre-sleep wake effects estimated using direct log-log fitting and *specparam* spectral parameterization. Both estimators revealed learning-related flattening of 1/f slopes over frontal and central regions relative to sham (log-log: $p_{perm} = 0.003$; *specparam*: $p_{perm} = 0.002$, cluster-based permutation tests). (b) Analogous Learning-minus-Sham maps shown for NREM stages 2-3. Both estimators demonstrated learning-related steepening of 1/f slopes in a frontocentral distribution (log-log: $p_{perm} = 0.049$; *specparam*: $p_{perm} = 0.002$, cluster-based permutation tests). White circles indicate electrodes included in significant cluster-corrected effects. Positive $t$ values indicate larger fitted slope values in learning relative to sham condition.

To determine whether the observed learning-related reorganization of aperiodic dynamics depended on the slope estimation procedure, we repeated the topographic analyses using both direct log-log fitting and *specparam* spectral parameterization fitted across 1-35 Hz using the average reference followed by the CSD transformation. In the log-log analysis, aperiodic slope was estimated by fitting a linear function to the log-transformed power spectrum, omitting frequency ranges dominated by prominent oscillatory activity from 8-16 Hz.

During pre-sleep wakefulness, both approaches identified a positive learning effect over the frontocentral regions, indicating learning-related spectral flattening. This effect reversed during subsequent NREM stages 2-3. Across both estimators, learning was associated with NREM spectral steepening over an overlapping frontocentral topography. The direction and state dependence of the effect were preserved, supporting the robustness of the wake-to-NREM inversion in learning-related aperiodic dynamics.

### 8 Robustness of 1/f slope effects to the analysis window

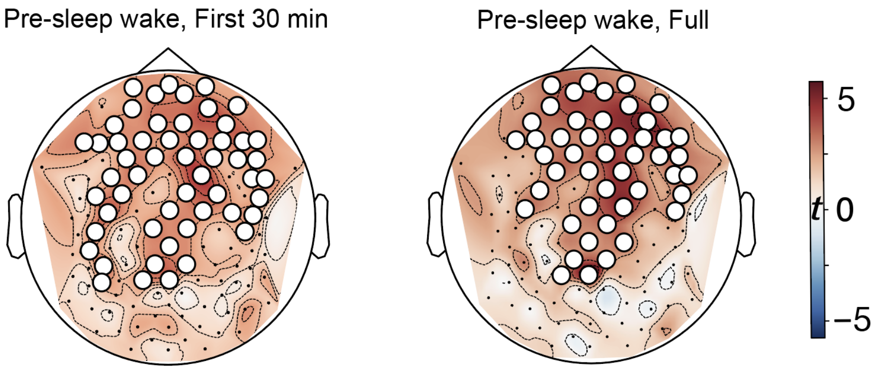


**Fig. S8.1 — The primary 30-min pre-sleep wake topography is preserved across the full wake interval.**

Notes. Learning-minus-Sham topographic $t$-statistic maps of pre-sleep wake aperiodic 1/f slope estimated using either the first 30 min of wakefulness (left) or the full available pre-sleep wake interval (right). White circles indicate electrodes belonging to significant sensor-space clusters.

The primary wake–NREM analyses restricted pre-sleep wakefulness to the first 30 min so that the waking estimate was defined over a temporal interval comparable to the first 30 min of NREM stages 2–3. We therefore repeated the pre-sleep wake topographic analysis using the full available wake interval to determine whether this temporal cap affected the learning-related spatial pattern (Fig. S8.1). The first-30-min and full-interval analyses showed closely corresponding topographies, with both revealing broad Learning-related flattening across the scalp that was strongest over the frontocentral region, with substantial overlap in the cluster-corrected electrodes. Therefore, the principal pre-sleep wake effect was not dependent on restricting the analysis to the first 30 min.

### 9 Electrode-level epoch-wise distribution of 1/f slopes

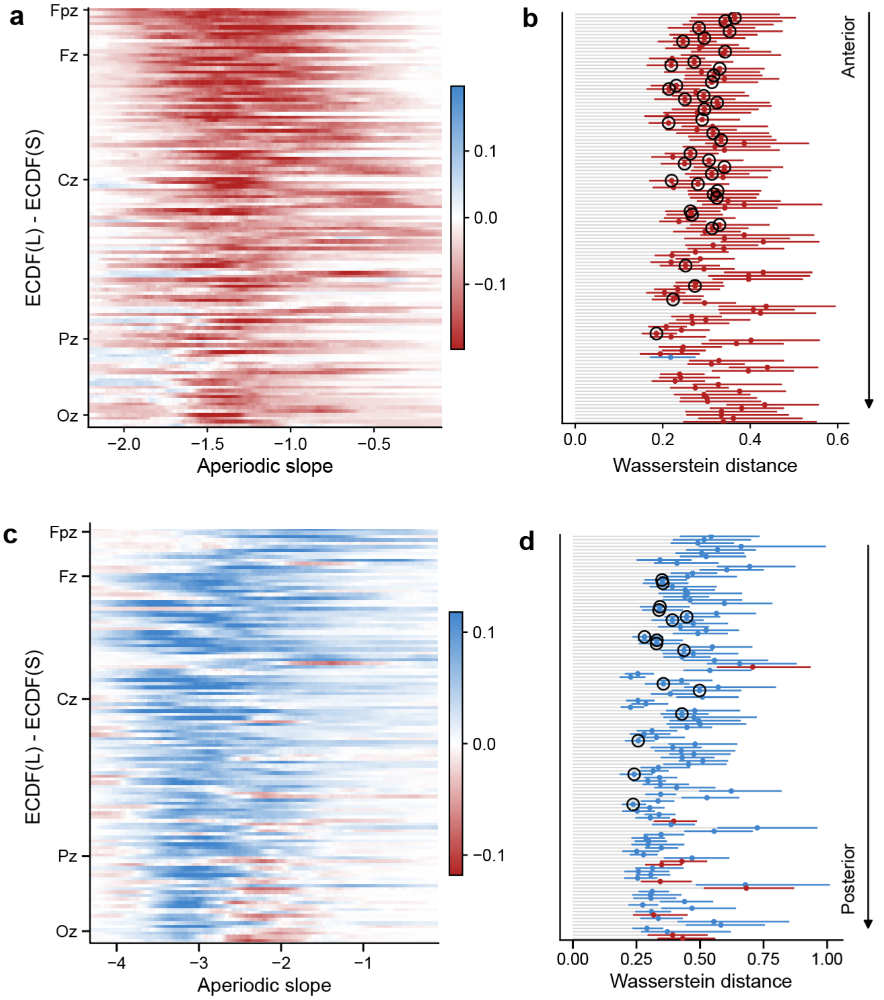


**Fig. S9.1 — Electrode-level distributional topology of aperiodic slopes across wake and NREM sleep.**

Notes. (a) Group-mean participant-level ECDF-difference heatmap for pre-sleep wake, ordered from anterior to posterior electrodes. Warmer colors indicate negative ECDF differences, where the Learning ECDF lies below the Sham ECDF, consistent with a relative increase in flatter, less negative slopes following learning. Cooler colors indicate positive ECDF differences, consistent with a redistribution toward steeper slopes. (b) Electrode-wise Wasserstein distance between Learning and Sham distributions during pre-sleep wake, providing a non-parametric quantification of the optimal transport cost required to transform the Sham 1/f slope distribution into the Learning distribution. Red points denote Learning shifts toward flatter slopes, whereas blue points denote Learning shifts toward steeper slopes. Horizontal error bars indicate 95% bias-corrected and accelerated (BCa) bootstrap confidence intervals across participants. Outlined black markers indicate electrodes where the mean participant-level signed area under the ECDF-difference curve (sAUC) differed significantly from zero in a two-sided one-sample sign-flip permutation test after FDR correction across electrodes. (c-d) Analogous to a-b, shown for the first 30 minutes of NREM stages 2-3, where learning was associated with a redistribution toward steeper 1/f slopes, reversing the wake state pattern.

ECDF-difference heatmaps revealed a scalp-wide redistribution toward flatter aperiodic slopes, with the largest Learning–Sham shifts over frontal and frontocentral regions (Fig. S9.1a). Electrode-wise Wasserstein distances quantified the magnitude of distributional separation and revealed a pronounced anterior pattern of aperiodic reorganization toward flatter slopes during wakefulness. Directional inference based on participant-level sAUC identified 40 predominantly frontal and frontocentral electrodes that remained significant after FDR correction ($p_{\text{perm}} = 0.001$–$0.002, q_{\text{FDR}} = 0.041$, sAUC; Fig. S9.1b). This distributional shift shows a reciprocal inversion during subsequent NREM sleep. Directional sAUC inference identified predominantly frontocentral electrodes that survived FDR correction ($p_{\text{perm}} = 0.0001$–$0.006, q_{\text{FDR}} = 0.006$–$0.048$, sAUC; Fig. S9.1c-d).

### 10 Cross-participant and cross-epoch generalization of aperiodic patterns

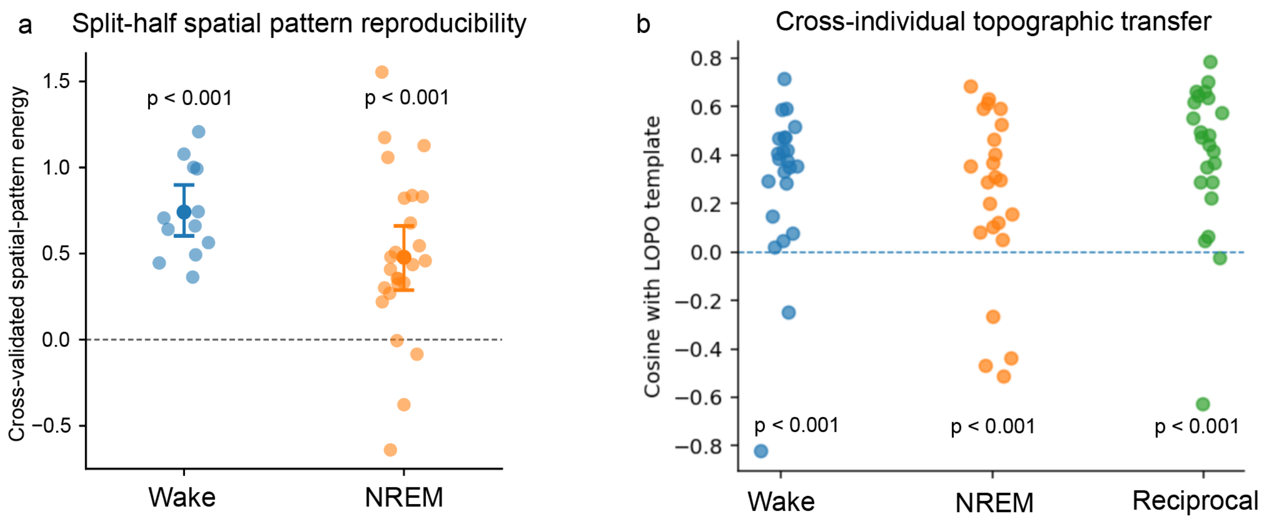


**Fig. S10.1 — Learning-related aperiodic spatial patterns are reproducible across time and generalize across participants.**

Notes. (a) Within-participant split-half reproducibility. Wake and early-NREM windows were divided into two non-overlapping subsets, and Learning-minus-Sham spatial patterns were estimated independently from each half. Cross-validated spatial-pattern energy quantified similarity between the independently estimated patterns. Analyses included 26 participants for Wake and 25 for NREM. Group significance was assessed using participant-level permutation tests, and points show individual participants with group means and 95% participant-bootstrap confidence intervals. Cross-validated spatial-pattern energy was significantly positive in both states ($P_{\text{perm}} < 0.001$). (b) Cross-participant generalization. For each held-out participant, Wake, NREM and reciprocal Wake–NREM spatial templates were estimated from the remaining participants using LOPO estimation and compared with the held-out participant's corresponding learning-effect topography using cosine similarity. Analyses included 26 participants for Wake, 25 for NREM and 23 for the reciprocal pattern. Group-level cosine similarity was tested against zero using participant-level permutation tests. Alignment was significantly positive for all three patterns (all $P_{\text{perm}} < 0.001$). Positive cosine similarity indicates alignment between the held-out participant's topography and the independently estimated template; dashed horizontal lines indicate zero similarity.

As a complementary sensitivity analysis to Fig. 5, we tested whether the spatial organization underlying the reciprocal wake–NREM effect was independently reproducible. First, we asked whether the Learning-minus-Sham topographies were stable across independent subsets of epochs within the same participant. Split-half validation showed significantly positive cross-validated spatial-pattern energy during both pre-sleep wake and early NREM (both $p_{\text{perm}} < 0.001$; Fig. S10.1a). Thus, the spatial structure of the learning-related effect was reproducible across time rather than depending on a particular subset of EEG windows.

We then asked whether these spatial patterns generalized across individuals. For each participant, wake, NREM, and reciprocal templates were estimated using only the remaining participants and compared with the held-out participant's corresponding topography. Held-out topographies showed significant positive alignment with the independently estimated templates for all three patterns (all $p_{\text{perm}} < 0.001$; Fig. S10.1b). Together, the split-half and leave-one-participant-out analyses show that the spatial organization contributing to the reciprocal effect was reproducible both across independent epochs within participants and across participants, supporting the stability of the spatial patterns used in the main state-space analysis.

#### 11 REM sleep aperiodic slope analysis

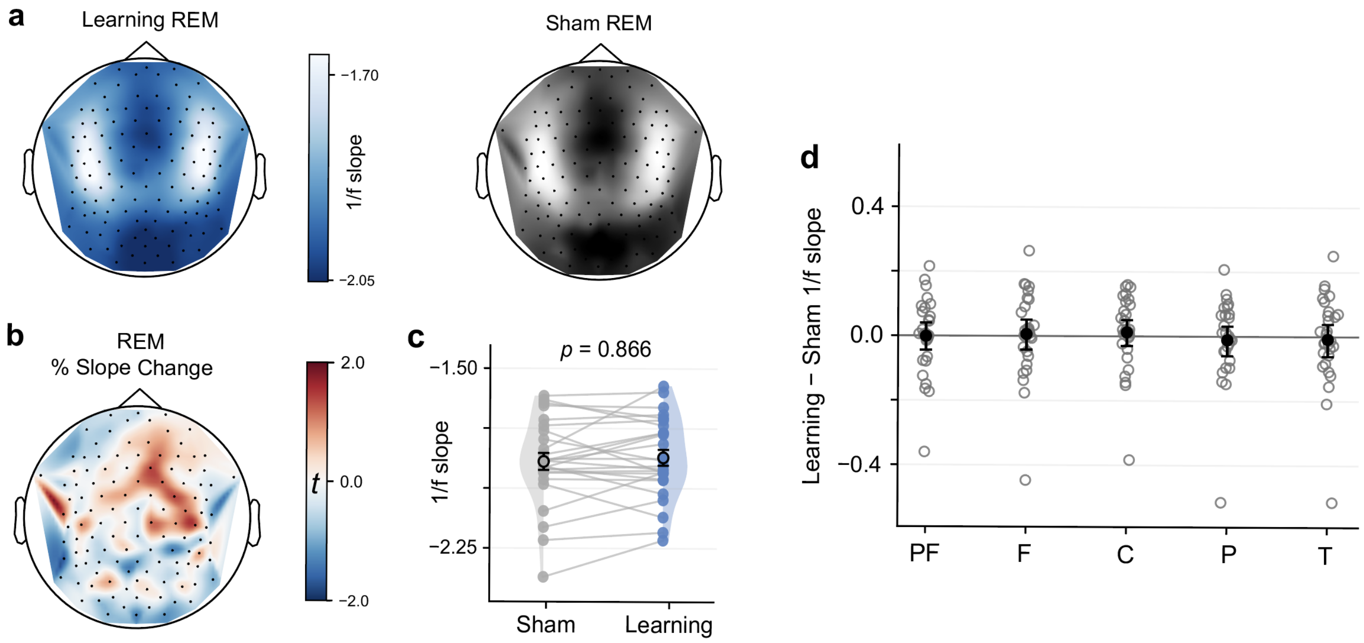


**Fig. S11.1 — REM sleep did not show learning-related modulation of aperiodic 1/f slope.**

Notes. (a) Topographic distribution of REM 1/f slope for learning and sham sessions. (b) Topographic $t$ map of slope change during REM sleep, computed as Learning minus Sham. (c) Participant-level paired plot of REM 1/f slope averaged across the frontal ROI, showing no significant difference between learning and sham sessions ($p_{perm} = 0.866$). (d) Learning-sham REM slope differences across predefined ROIs. PF, prefrontal; F, frontal; C, central; P, parietal; T, temporal.

To assess whether learning-related changes in aperiodic dynamics generalized across sleep states and whether REM also shows a learning-induced renormalization, we examined 1/f slope during REM sleep. In contrast to the frontocentral NREM steepening observed after learning, REM sleep showed no reliable learning-related change in 1/f slope. Topographic maps of REM slope change did not reveal a consistent spatial pattern, and participant-level ROI estimates were comparable between learning and sham sessions. The absence of a REM effect supports the state specificity of the learning-related aperiodic reorganization observed in NREM sleep, rather than reflecting a general sleep-stage difference between learning and sham nights.

#### 12 REM aperiodic slope and memory association

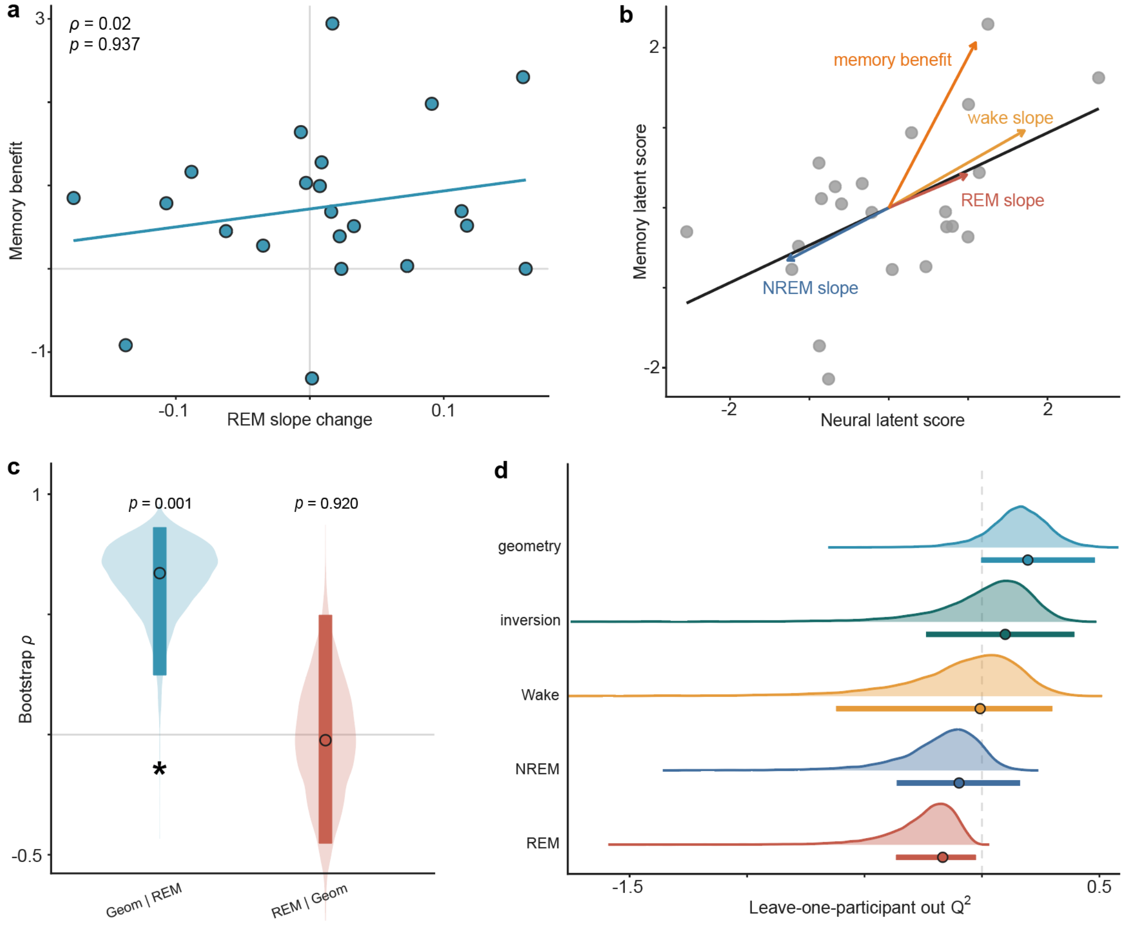


**Fig. S12.1 — REM aperiodic slope does not account for learning-related memory change.**

Notes. (a) REM Learning-sham 1/f slope differences were not associated with sleep-specific memory benefit ($\rho = 0.019$, $P_{\text{perm}} = 0.937$). (b) Partial least squares correlation analysis of Wake, NREM and REM aperiodic slope with memory benefit. Wake and NREM slopes projected in opposing directions on the neural latent axis, whereas the REM projection was smaller and Wake-aligned. (c) Cross-state geometry remained strongly associated with memory benefit after controlling for REM slope ($\rho = 0.671$, 95% BCa CI, 0.247–0.862, $P_{\text{perm}} = 0.001$), whereas REM slope showed no association after controlling for geometry ($\rho = -0.024$, 95% BCa CI, -0.453–0.498, $P_{\text{perm}} = 0.920$). Violins show participant-bootstrap distributions. (d) Leave-one-participant-out prediction of memory benefit using the same models as in the main analysis, with REM added as a control. REM did not show significant held-out prediction for memory benefit ($Q^2 = -0.167$, $q_{\text{FDR}} = 0.526$). Ridges show participant-bootstrap distributions. For (c) and (d), thick bars show 95% BCa CIs and circles show observed values.

We further tested whether REM aperiodic slope contributed to the neural–behavioral relationship identified across Wake and NREM. REM Learning-minus-Sham slope was not associated with sleep-specific memory benefit (Fig. S12.1a). A three-state PLSC showed opposing Wake and NREM slope projections, with REM exhibiting a weaker, Wake-aligned projection (Fig. S12.1b). The relationship between cross-state geometry and memory benefit remained strong after controlling for REM slope, whereas REM slope still showed no association after controlling for geometry (Fig. S12.1c). Held-out prediction showed the same pattern: cross-state geometry and static inversion showed significant positive $Q^2$ after FDR correction, whereas REM slope did not predict memory benefit out of sample (Fig. S12.1d). Thus, the neural–memory relationship was specifically captured by coordinated Wake–NREM organization rather than by learning-related aperiodic slope changes extending into REM sleep.

#### 13 Neural associations with retrieval-latency change

| Neural measure | $N$ | $\rho$ | $p_{\text{perm}}$ | $q_{\text{FDR}}$ |
|---|---|---|---|---|
| Wake 1/f slope | 22 | 0.252 | 0.260 | 0.458 |
| NREM 1/f slope | 22 | -0.238 | 0.286 | 0.458 |
| Wake–NREM inversion | 22 | 0.355 | 0.101 | 0.406 |
| Cross-state geometry | 22 | 0.356 | 0.102 | 0.406 |
| Common mode | 22 | -0.126 | 0.572 | 0.572 |
| Pre-sleep theta | 22 | 0.308 | 0.167 | 0.444 |
| NREM SWA | 22 | -0.147 | 0.515 | 0.572 |
| NREM sigma | 22 | 0.130 | 0.562 | 0.572 |

**Table S13.1 — Learning-related neural measures are not associated with sleep-specific changes in retrieval latency.**

Notes. Sleep-specific retrieval-latency change was defined as the Sleep-minus-Wake difference in delayed-minus-immediate log mean RT on correct trials. Spearman correlations quantify associations between this behavioral measure and participant-level Learning-minus-Sham neural measures. $p$ values were obtained from two-sided permutation tests. $q$ values were calculated using Benjamini–Hochberg correction across neural measures.

Given that retrieval latency changed across the retention interval, we next asked whether individual differences in this behavioral change were associated with the same learning-related neural measures examined for memory accuracy. None of the neural measures was significantly associated with retrieval-latency change (Table S13.1; all $p_{\text{perm}} > 0.10$, all $q_{\text{FDR}} > 0.40$). Therefore, the neural–behavioral relationships observed for memory accuracy did not generalize to individual differences in response speed.
